# Foveation as a self-supervision signal: Blurring beats blanking for learning robust visual representations

**DOI:** 10.64898/2026.09.26.754698

**Authors:** Akihito Maruya, Hossein Adeli, Tian Zeng, Nikolaus Kriegeskorte, Ning Qian

**Affiliations:** Zuckerman Institute Columbia University; Data Science Institute Columbia University

## Abstract

The primate retina is strikingly non-uniform, with receptor density falling off sharply from fovea to periphery. Every eye-movement brings a previously low-acuity peripheral area to high-acuity foveal processing. We hypothesize that this process provides a natural self-supervision signal, enabling representation learning by predicting fine details in the periphery. We implement this in a SimMIM-style vision transformer, replacing its blank masks (ViT-Blank) with a uniform blur mask (ViT-Blur) and further with retinal masks in which a foveal patch is randomly selected, and all other patches are progressively blurred based on eccentricity from the foveal patch (ViT-Retina). ViTs were pre-trained to reconstruct the full-resolution image from the corrupted input and then fine-tuned on clean images for classification. ViT-Blur and ViT-Retina outperform ViT-Blank on the pre-training reconstruction task while controlling for retained information from the input across corruption types. Critically, they also outperform ViT-Blank on downstream classification, showing that they learn representations that generalize better. The ViT-Blur and ViT-Retina are also more robust in impoverishment conditions such as higher mask ratios, fewer pre-training epochs, smaller training sets, and test time image corruptions. Using additive band-limited noise, we discovered that ViT-Blank relies on higher spatial frequencies compared to ViT-Blur and ViT-Retina, and we show this to be the likely cause of poorer generalization: the centroid of a model’s spatial frequency tuning is negatively correlated with its classification accuracy (*r* = −0.73). We further show that a closed-form linear-ridge encoder derived from the same reconstruction objective reproduces this tuning (*r* = 0.88). Overall, our experiments show the promise of a biologically-inspired self-supervision objective for learning robust visual representations.

## 1 Introduction

Photoreceptor density in the primate retina falls off significantly with eccentricity (Curcio et al., 1990; Wandell, 1995; Watson, 2014). Every saccade delivers a different sampling of the visual world (Loschky et al., 2005), yet the visual system assembles the complementary information across fixations into a coherent percept. We hypothesize a continuous cycle of prediction and confirmation: the visual system anticipates peripheral structure and receives foveal feedback after each saccade. Over time, the cycle of prediction and correction provides a natural self-supervision signal that may drive the acquisition of highly robust and generalizable visual representations.

Masked image modeling (MIM), predicting masked patches from the visible ones, has been a very effective objective for learning robust visual representation that transfers to a range of downstream tasks (He et al., 2022; Xie et al., 2022). Common masking methods in the literature replace the patch tokens with *blank* (Xie et al., 2022) or drop them entirely (He et al., 2022). The primate peripheral vision however uses a different form of corruption: the progressive blurring of the image with increasing eccentricity contains bandwidth-limited signal across the entire visual field. Here we take inspiration from foveated vision and explore biologically inspired masking in MIM. During pre-training we divide each image into patches and corrupt a randomly chosen subset of them. The model is trained to reconstruct the original image.

We study three families of corruption. **Blank** masking fills the selected patches with mid-gray (ViT-Blank). **Blur** masking replaces the blank masking with uniform band-limited blur (ViT-Blur). **Retinal** masking filters every patch at a resolution that decays with increasing its eccentricity from a randomly drawn fixation patch (ViT-Retina). We then evaluate the quality of the learned representation by fine-tuning each pre-trained model on clean images for classification. We ask: **Q1** Can ViT-Blur and ViT-Retina reconstruct full-resolution detail better than ViT-Blank when retained information are matched across corruption types, and is there a benefit to the more biologically inspired blurring in ViT-Retina? **Q2** Does the corruption type have a meaningful impact on learning transferable and robust representations? **Q3** What mechanisms can explain the potential differences among them?

### Contributions

(1) Using the MIM framework, we evaluate the effect of three main corruption types with different degrees of biological inspiration on both the reconstruction performance and representation transfer to a classification task. Across CIFAR-10, ImageNet-100 and ImageNet-1k, we observe that ViT-Blur and ViT-Retina outperform ViT-Blank on both metrics. Furthermore, they are more robust to impoverishment conditions such as higher pre-training mask ratios, fewer pre-training epochs, smaller training sets, and test time image corruptions. (2) We show that a probe using spatial-frequency (SF)-band-limited noise explains the classification gap: ViT-Blank relies on high-SF under impoverishment conditions, while ViT-Blur and -Retina models do not, and the centroid of a model’s SF tuning is negatively correlated with its classification accuracy (r = −0.73). (3) We solve a closed form expression for a linear encoder with the same pre-training objective and reproduce the the ViT’s SF tuning (*r* = 0.88) using the same band-limited probe as above. We further examine the mechanism in this simpler model and show that the biologically motivated masking emphasizes the mode carrying more class signal.

## 2 Related work

### Masked image modeling (MIM)

MIM is a powerful pre-training objective for images (He et al., 2022; Xie et al., 2022) and video (Feichtenhofer et al., 2022; Tong et al., 2022; Wang et al., 2023), underpinning foundation models such as Segment Anything (Kirillov et al., 2023) and InternVideo (Wang et al., 2022), and supporting encoders of human visual responses (Adeli et al., 2026b; Hwang et al., 2025; Adeli et al., 2026a). Built on ViTs (Dosovitskiy et al., 2020), MAE (He et al., 2022) and SimMIM (Xie et al., 2022) reconstruct missing content from partial observations. MAE (He et al., 2022) drops the masked tokens entirely and encodes only the visible ones, then uses a decoder to insert mask tokens and reconstruct the masked image patches. SimMIM (Xie et al., 2022) instead keeps the full grid, masks at the token level rather than dropping them, and reconstructs through a single linear decoder. Random patch masking approach is inherited from language modeling, where the masked-token objective was introduced for Transformers (Vaswani et al., 2017; Devlin et al., 2019), but how the masking is done matters. Hinojosa et al. (2024) generates data-independent binary masks by filtering noise in different frequency bands and finds that this corruption method leads to better transfer of the learned representation compared to random masking of standard MAE, which suggests that the frequency content of the mask shapes what is learned.

### Theoretical accounts of MIM

Recent theoretical work has sought to explain MAE’s empirical success: Kong et al. (2023) formalize MAE via a hierarchical latent-variable model and show how mask ratio and patch size determine which latent variables are recovered. Zhang et al. (2022) show that MAE implicitly performs contrastive learning. Reconstructing one target from two different views pulls the representations of those views together, as a positive pair in the contrastive learning would do. Their derived mask ratio closely matches the empirically optimal one. A complementary line of work analyzes self-supervised methods through linear autoencoders, starting from the classical results that an autoencoder is closely related to principal component analysis (PCA) of the input data (Bourlard & Kamp, 1988; Baldi & Hornik, 1989; Plaut, 2018). Bisulco et al. (2025) extend this analysis to MAE, deriving a closed-form linear-MAE encoder by marginalizing the binary mask in expectation and showing that mask ratio and patch size shape which spatial correlations the encoder learns; they then bridge linear and non-linear MAE through a first-order Taylor expansion of the trained encoder, whose Jacobian inherits spectral structure from the closed-form optimum. Because our tested corruption is non-binary (low-pass blur), this mask-marginalization machinery does not apply directly; we instead take the corrupted-input matrix *X*_*c*_ as the observable and apply the SVD to it directly, which lets us treat *corruption type* as a mechanistic axis alongside the mask ratio, pre-training budget and training-set size (§6).

### Foveation, spatial frequency, and biological vision

A parallel line of work examines how deep artificial vision systems exploit SF and blur information. Foveated perceptual systems trained with peripheral blur develop emergent properties that match aspects of human vision (Deza & Konkle, 2020; Wang & Cottrell, 2017), and Pramod et al. (2022) show that the natural retinal blur profile is, in fact, *optimal* for object recognition — a direct motivation for replacing blank masks with biologically plausible blur. Vogelsang et al. (2018) shows that a network trained first on blurred faces then on high-resolution ones develops larger receptive fields, supporting configural processing, whereas a network trained with reversed order does not. Visual processing is often coarse-to-fine Schyns & Oliva (1994); Chen & Qian (2004) with low-spatial-frequency information guiding the processing of fine details. Wang et al. (2020) and Yin et al. (2019) show that CNNs disproportionately rely on high-spatial-frequency content that humans largely ignore, and Li et al. (2023) show that robust object-recognition models instead lean on low-frequency components, suggesting that coarse spatial structure supports stability against perturbations. Geirhos et al. (2018b) systematically compare humans and CNNs on noisy and blurred images and show that the human–network gap widens precisely under such corruptions, motivating training regimes that close it. A related thread argues that the gap is partly a shape-vs-texture bias: ImageNet-trained CNNs are texture-biased while humans are shape-biased, and increasing shape bias improves accuracy and robustness (Geirhos et al., 2018a; Hermann et al., 2020). In the context of modeling human vision, Jang & Tong (2024) demonstrate that incorporating blur robustness into CNNs improves alignment with human perceptual responses. In the previous work, the biologically motivated degraded input is either an augmentation for the input images in supervised classification or a front end built into the architecture. We instead integrate biologically motivated masking into the MIM framework.

## 3 Method

### 3.1 Pipeline and corruption types

The pipeline has two stages: (1) masked-reconstruction pre-training with a SimMIM-style encoder and one-layer decoder (Xie et al., 2022), reconstructing the *full-resolution* patches from the corrupted input under an *L*_1_ loss; and (2) fine-tuning on *clean* images for classification via the `[CLS]`token.

We adopt a modified version of the SimMIM approach and apply the corruption to the image rather than the tokens. This allows us to compare blank, blur, and retinal masking fairly as all the corruptions modify the image content themselves. For example, the **ViT-Blank** fills a masked patch with mid-gray. We also include token masking as used in SimMIM (Xie et al., 2022) as a control and provide the results for this baseline.

We use four blur masking conditions. **ViT-Blur (strong)** applies a 2-D isotropic Gaussian low-pass (*σ* = 5.09 px on CIFAR-10, 35.65 px on ImageNet). **ViT-Blur (strong)** + **3 cyc/image** and **ViT-Blur (strong)** + **2 cyc/image** apply the same Gaussian and additionally set to zero every Fourier coefficient above 3 and 2 cyc/image, respectively. **ViT-Blur (mild)** applies a narrower Gaussian (*σ* = 1.27 px on CIFAR-10, 8.91 px on ImageNet) and sets Fourier coefficients to zero for frequencies greater than 8 cyc/image, so it both blurs less and keeps a wider passband than the other three. The cutoff conditions are motivated by the retinal Nyquist limit at progressively more peripheral eccentricities (Curcio et al., 1990; Watson, 2014): SF content above that limit is absent from the retinal signal, whereas a Gaussian blur only attenuates it and is invertible in principle.

**ViT-Retina** uses a masking whose resolution falls with distance from a fixation. One patch per image is designated as the fovea, drawn uniformly at random, and is left unblurred; every other patch is low-pass filtered at a cutoff that falls with its distance from that fixation, following the human retinal acuity falloff (Geisler & Perry, 1998). The field of view (FOV) sets the scale: at FOV 20° the diagonal corners of the image subtend 20° of visual field, so a larger field of view makes the frequency cutoff decay faster across the image resulting in a more blurred periphery (App. A.2).

A blank mask throws the content of a patch away, while blur and retinal masks keep its low spatial frequencies. In order to have a fair reconstruction accuracy comparison, we matched the retained information in the images following each corruption, formalized as **retained degrees of freedom** (DOF), across the corruption types (App. A.3). For blank masking we count the number of pixels of the visible patches as the masked patches carry no information. For uniform blur masking we count those same visible pixels and add the number of non-zero Fourier coefficients the band-limited filter leaves. For retinal masking we count the pixels of the sharp foveal patch plus the non-zero Fourier coefficients of each eccentricity ring, weighted by the area that ring covers, averaged over fixations.

### 3.2 Architecture, data, and training regimes

We use a ViT with a SimMIM-style 1 layer decoder (Xie et al., 2022): depth 6 with embedding dimension 384 on CIFAR-10 (Krizhevsky et al., 2009), and ViT-B (depth 12, dimension 768) on both ImageNet100 and 1k (Russakovsky et al., 2015). All the parameters are in App. A.1. **Masks are re-sampled every forward pass**, so fewer epochs mean fewer distinct mask arrangements per image. We denote epoch budgets as *pre-training* + *fine-tuning*: e.g. 200+100 means 200 pre-training epochs followed by 100 fine-tuning epochs.

The mask ratio is the fraction of image patches that are corrupted, so a higher mask ratio leaves less of the image intact and makes the reconstruction harder (Figs. 1 and S2). On CIFAR-10, we use mask ratios of {0.55, 0.65, 0.75, 0.85, 0.95} and FOVs of {2.5, 5, 10, 20, 40, 80, 160} ° at 200+100 epochs on 40K training images; we also run 150+100, 100+100 and 50+100 epochs to determine how efficiently the models pre-trained under each corruption type use the pre-training budget, and 200+100 epochs at 30K, 20K, and 10K training samples, to see how robust they are to data scarcity. On ImageNet-100, we use mask ratios {0.65, 0.95} and FOV {20, 40, 160} ° at 800+100 epochs; we also run 100+100 epochs for reduced pre-training, and 0.2 of the training set for reduced training samples. On ImageNet-1k, we use mask ratio {0.65, 0.95}, FOV {20, 40, 160}° at 100+100 epochs.

**Figure 1:**
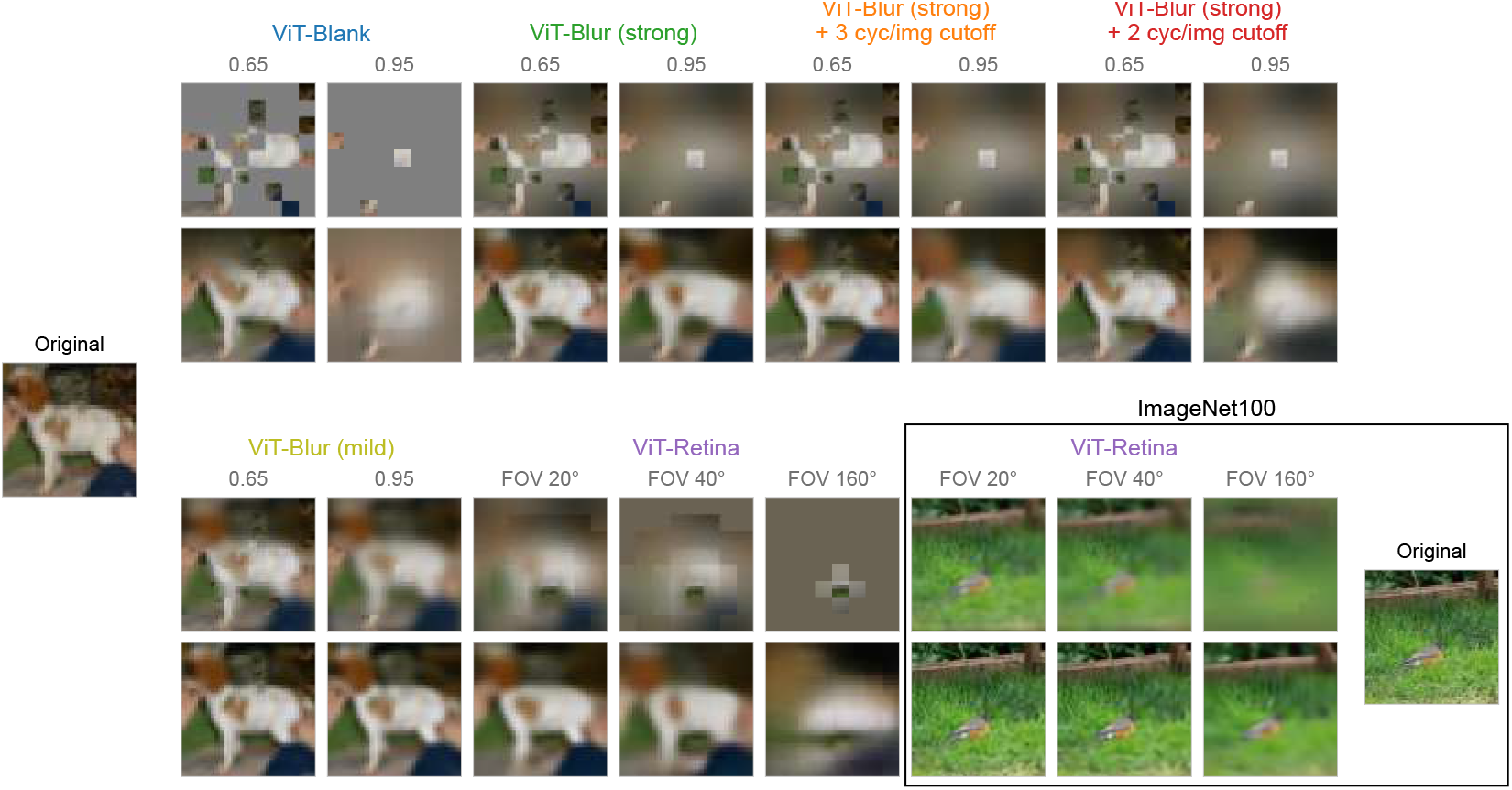
Pre-training inputs and reconstructions. One CIFAR-10 image under each corruption (top row of each block) and the corresponding model’s reconstruction (bottom): ViT-Blank and the ViT-Blur models at mask ratios 0.65 and 0.95, and ViT-Retina at FOV 20, 40 and 160°. Boxed, right: ViT-Retina on ImageNet-100.

### 3.3 Evaluation

All metrics are computed on held-out test data that is never used during pre-training or fine-tuning. We measure pre-training reconstruction quality as the *L*_1_ distance between the decoder output and the clean image, evaluated on test images corrupted as in pre-training. Examples are shown in Fig. 1 (CIFAR-10 and ImageNet100) and Fig. S2 (ImageNet), and quantitative errors against mask ratio in Fig. S3 and against retained DOF in Fig. S5.

We then apply four downstream evaluations to the fine-tuned classifier. First, the *classification probe* measures test accuracy on clean images, and we plot that accuracy against the pre-training mask ratio and the FOV in Figs. 2 and S8, and against the retained DOF of the pre-training input in Fig. S9. Second, the *noise-robustness probe* adds Gaussian noise (*µ* = 0, *σ* = 0.05 on [0, 1] pixel values) to each test image at evaluation time and measures accuracy on those noisy images; Figure 2D plots it against the pre-trained mask ratio and the field of view on CIFAR-10, and Fig. S10 plots it on every dataset, against both the pre-trained mask ratio and the retained DOF. Third, *corruption probe* measures accuracy on CIFAR-10C, ImageNet-100-C, and ImageNet-1k-C, which applies 15 image corruptions at 5 severities (Hendrycks & Dietterich, 2019), and the accuracy is plotted against the pre-trained mask ratio in Fig. S11 and against the retained DOF in Fig. S12. Fourth, the *SF-band-limited noise probe* adds noise built by multiplying the same white noise used in the noise-robustness probe by a Gaussian envelope (*σ*_*f*_ = 0.52 cyc/image) in Fourier space, centered at a chosen spatial frequency between 1 and 15 cyc/image, and plots the resulting accuracy drop (clean minus noisy) against the central frequency of the Gaussian envelope on CIFAR-10 (Fig. 3). Noise is *never* applied during pre-training or fine-tuning.

**Figure 2:**
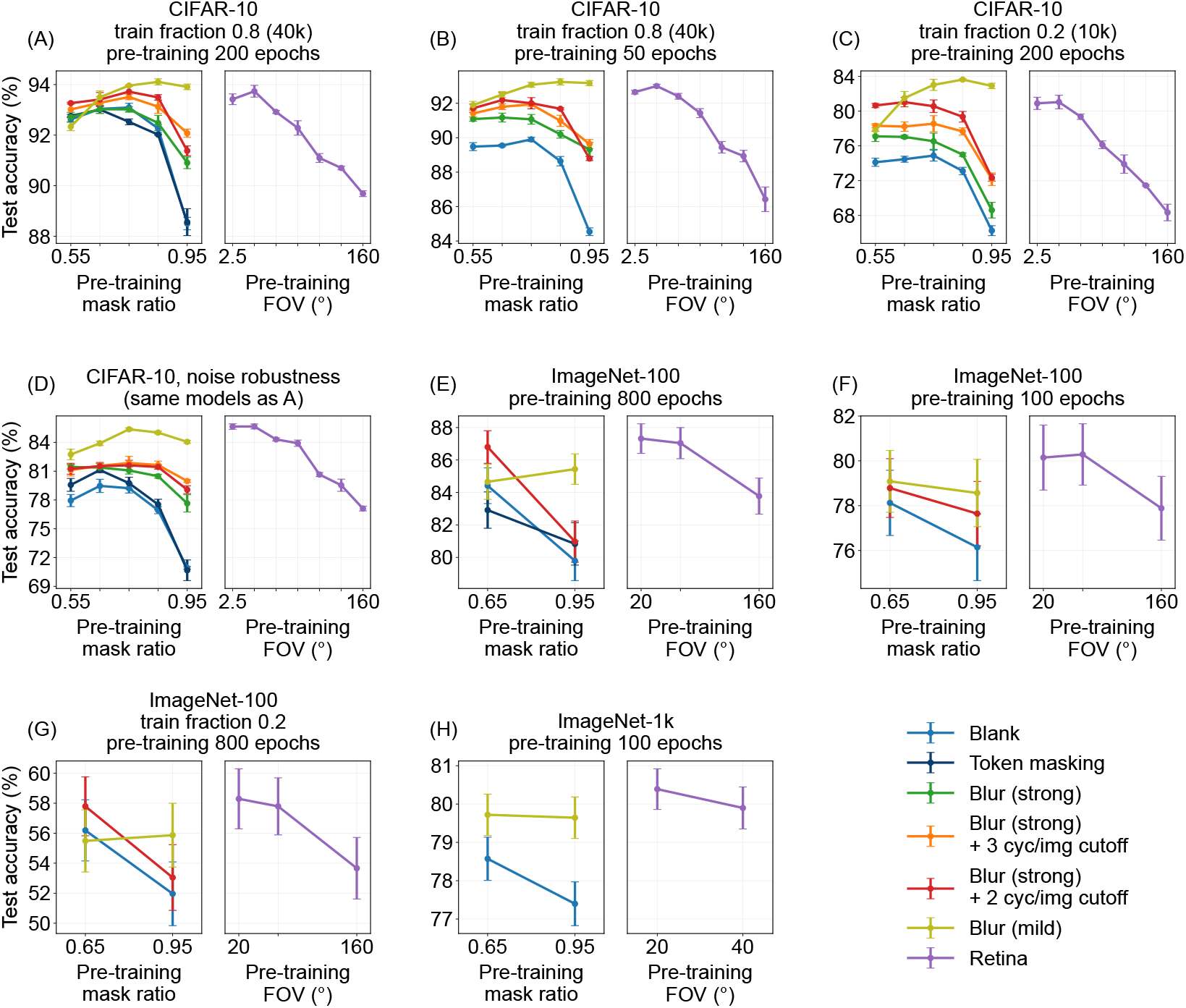
Classification accuracy after fine-tuning. **(A)** CIFAR-10 at 200+100 epochs on 40K images; **(B)** a short pre-training budget (50+100 epochs); **(C)** a reduced training set (10K images); **(D)** the models of A under Gaussian pixel noise (*σ* = 0.05). **(E)** ImageNet-100 at 800+100 epochs on the full training set; **(F)** a short pre-training budget (100+100 epochs); **(G)** a reduced training set (a fifth of the images); **(H)** ImageNet-1k at 100+100 epochs. Panels A–D average three seeds; E–H are single runs.

**Figure 3:**
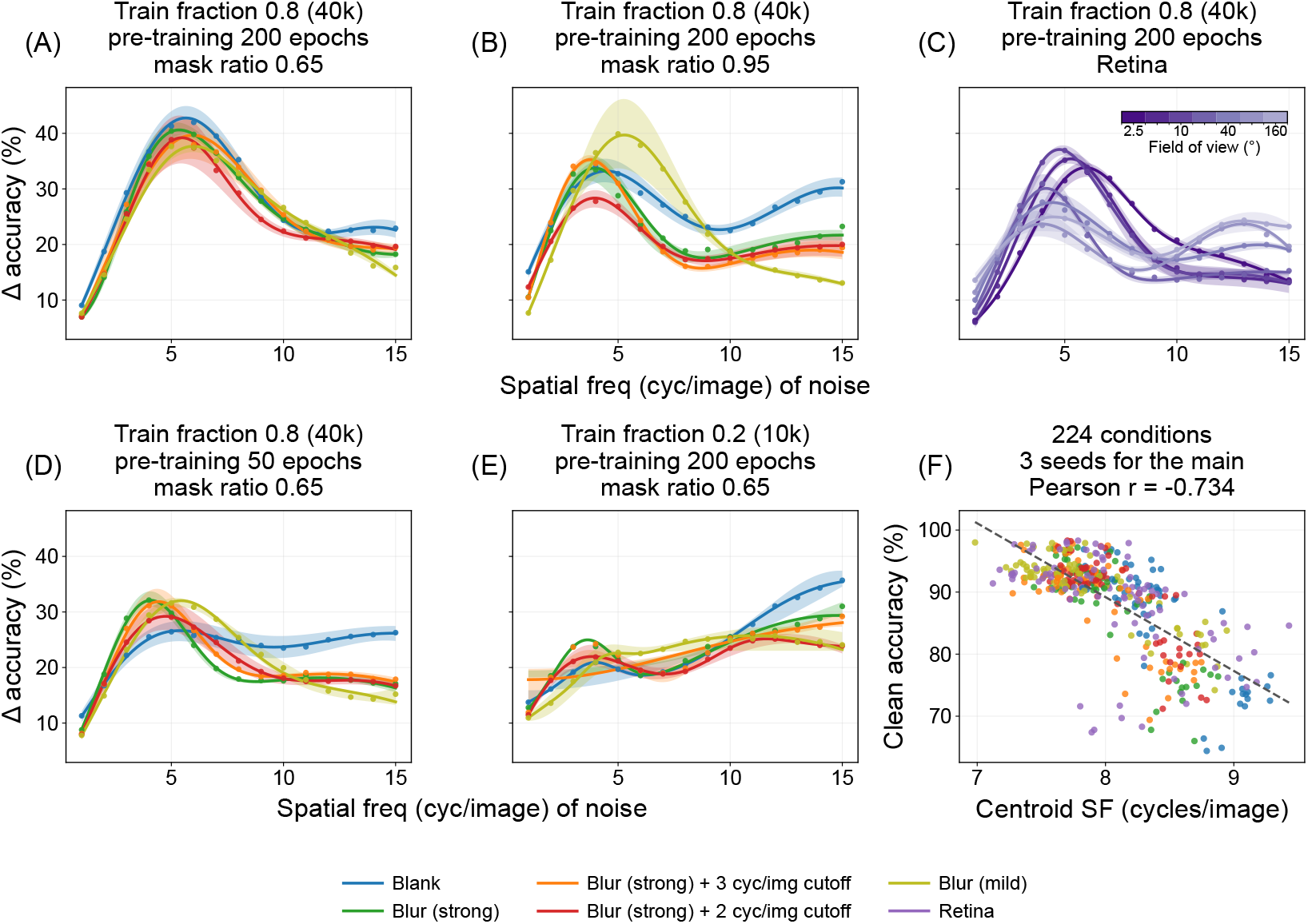
Spatial-frequency tuning of the fine-tuned models. Accuracy drop under band-limited noise against the band’s centre frequency; higher means the model relies more on that band. **(A, B)** full condition at pre-training mask ratio 0.65 and 0.95; **(C)** ViT-Retina (2.5–160°); **(D)** 50-epoch budget and **(E)** train fraction 0.2, both at mask ratio 0.65. **(F)** Clean accuracy against tuning-curve centroid for all 224 models, one point per seed (*n* = 416), with the least-squares line (*r* = −0.73).

For CIFAR-10 we use an 80*/*20 split of the 50K training images (40K train, 10K test), and the reduced-sample conditions subsample 30K, 20K or 10K from that training set; the 10K test set is held fixed throughout. For ImageNet we use the standard train/validation split, with the ImageNet-100 reduced-sample condition subsampling 0.2 of the training set and every model evaluated on the same validation set.

## 4 ViT-Blur/Retina reconstructs fine detail and transfers better

### Reconstruction (Q1)

ViT-Blur achieves consistently lower *L*_1_ error than ViT-Blank (as well as token masking) across every mask ratio and every impoverishment conditions (CIFAR-10, ImageNet-100, and ImageNet-1k: Fig. S3), and the blur-vs-blank gap widens with increasing mask ratio. This is also visible qualitatively: at moderate mask ratio 0.65 all corruption types yield coherent reconstructions on CIFAR-10 (Fig. 1) and ImageNet (Fig. S2), but at mask 0.95 ViT-Blank produces fragmented outputs that fail to recover global shape while ViT-Blur and ViT-Retina still reconstruct coherent objects and scenes. Within the strong blur family, reconstruction error increases monotonically as the cutoff frequency is lowered (Gaussian blur (green) *<* 3 cyc/image (orange) *<* 2 cyc/image (red)), reflecting the progressive loss of SF information in the input.

Because blur and retinal masks retain a patch’s low SFs, the fair comparison is at nearly matched DOF (§3.1). Re-plotting every condition against retained DOF (Fig. S5; accounting in App. A.3), reconstruction *L*_1_ error follows ViT-Retina *<* ViT-Blur *<* ViT-Blank on every dataset. This answers **Q1**: a ViT reconstructs detail better from band-limited input than from blanked input at matched DOF, and the more biologically motivated ViT-Retina reconstructs better than ViT-Blur.

A further concern is that the blur-trained ViT is simply learning to invert the blur kernel. We rule this out at mask ratio 1.0, where the whole image is blurred and no sharp patch is left to infer from: a Wiener deconvolution given the ground-truth kernel is still substantially worse than the ViT, which never sees the kernel (Fig. S6). The encoder has therefore learned image structure beyond kernel inversion. Below we show whether the learned structure is actually useful for downstream classification tasks.

### Classification (Q2)

Figure 2 plots classification accuracy after fine-tuning against the pre-training mask ratio and FOV. ViT-Blur and ViT-Retina outperform ViT-Blank (as well as token masking) in classification accuracy for every dataset. All impoverishment conditions lower accuracy for every model, but ViT-Blur and ViT-Retina are more robust to them than ViT-Blank. In the full condition, which uses the largest number of pre-training epochs and full training samples for each dataset, the best ViT-Blur leads ViT-Blank by 0.7% at pre-training mask ratio of 0.55 and by 5.4% at pre-training mask ratio of 0.95. That lead at pre-training mask ratio of 0.95 grows to 8.7% under short budget (Fig. 2B) and to 16.6% in train fraction 0.2 (Fig. 2C). For ViT-Retina, accuracy rises as the pre-training FOV narrows (Fig. 2A), and the peak stays at 5° under the short budget and reduced data as well. Within the blur family, ViT-Blur (mild) is the best uniform blur at every pre-training mask ratio of 0.65 and above. Within ViT-Blur (strong) family, imposing a cutoff in pre-training masking helps in classification accuracy (Gaussian blur (green) *<* 3 cyc/image (orange) ≈2 cyc/image (red)), the reverse of the reconstruction ordering.

On ImageNet-100 and ImageNet-1k, the same pattern is observed. In the full condition (Fig. 2E), ViT-Blur beats ViT-Blank (and token masking) at both mask ratios: ViT-Blur (strong + 2 cyc/image) leads ViT-Blank by 2.4% at pre-training mask ratio of 0.65, and ViT-Blur (mild) by 5.6% at pre-training mask ratio of 0.95. The fine-tuning accuracy of ViT-Retina is ordered by the pre-training FOV (FOV 20° best at 87.3%, 40° at 87.0%, 160° at 83.8%), with FOV 20° being above all ViT-Blur and ViT-Blank on classification accuracy. The ViT-Blur’s and ViT-Retina’s advantages over ViT-Blank hold for shorter pre-training budget (Fig. 2F) and reduced data (Fig. 2G). The ordering holds as we scale to the ImageNet-1k (Fig. 2H): at pre-training mask ratio of 0.95 ViT-Blur (mild) reaches 79.6% against ViT-Blank’s 77.4%, and ViT-Retina at FOV 20° outperforms them at 80.4% accuracy. The full condition results for all three datasets are summarized in Table 1, and every impoverishment condition in Table 3.

**Table 1:** Accuracy in the full condition. Clean, Gaussian-noise (noise) and common-corruption (-C) top-1 accuracy for the best model of each family. Bold marks the best of the three in each column. Every impoverishment condition is given in Table 3.

| Dataset | ViT-Blank |  |  | ViT-Blur |  |  | ViT-Retina |  |  |
| --- | --- | --- | --- | --- | --- | --- | --- | --- | --- |
|  | clean | noise | -C | clean | noise | -C | clean | noise | -C |
| CIFAR-10 (200+100 ep) | 93.1 | 79.2 | 79.2 | **94.1** | 85.0 | **82.1** | 93.7 | **85.7** | 81.4 |
| ImageNet-100 (800+100 ep) | 84.4 | 79.6 | 52.9 | 86.8 | 82.1 | 58.0 | **87.3** | **84.8** | **62.7** |
| ImageNet-1k (100+100 ep) | 78.6 | 75.7 | 52.4 | 79.7 | 77.3 | 56.4 | **80.4** | **78.4** | **58.3** |

To compare ViT-Blank and ViT-Blur with ViT-Retina on a common axis, we re-plot the classification accuracy against retained DOF (Fig. S9). On CIFAR-10 the pattern is common across the pre-training corruptions: starting from the strongly masked end the accuracy rises with increasing DOF, and peaks around 150–350 retained DOF, and then falls as the DOF continues increasing towards the lightly masked end. In the full condition, ViT-Blur (mild) performs the best at nearly equal DOF, followed by ViT-Retina, then the ViT-Blur (strong) family, with ViT-Blank coming last. The same ordering holds under every impoverishment condition, and the gap from ViT-Retina and the ViT-Blur (strong) family down to ViT-Blank widens as the conditions are further impoverished. On ImageNet-100 and ImageNet-1k, ViT-Retina performs the best at nearly matched DOF, then ViT-Blur, and then ViT-Blank (Fig. S9).

### Robustness

Under Gaussian noise the gap between ViT-Blur/-Retina and ViT-Blank becomes wider (see Table 1 for best model performance for all pre-training schema, test datasets and test time images corruptions). For example, on CIFAR-10, at pre-trained mask ratio 0.95, ViT-Blur (mild) leads ViT-Blank by 13.2% under noise against 5.4% on clean images (Fig. 2D). On ImageNet-100 the lead of ViT-Retina at FOV 20° over ViT-Blank at mask ratio 0.65 grows from 2.9% on clean images to 5.2% under noise, and on ImageNet-1k from 1.8% to 2.7%. The advantage also becomes larger at matched information on the DOF axis, where ViT-Retina is the top performing model (Fig. S10D-F). The same holds on CIFAR-10-C, ImageNet-100-C and ImageNet-1k-C (Figs. S11 and S12). On CIFAR-10-C ViT-Retina and ViT-Blur (mild) sit highest at every DOF and ViT-Blank lowest. On both ImageNet sets, in the full condition, the lead for ViT-Retina at FOV 20° over ViT-Blank at mask ratio 0.95 widens once the images are corrupted during test. On ImageNet-100 it grows from 7.5 points on clean images to 12.1. On ImageNet-1k it grows from 3.0 points to 5.5. Together these results answer **Q2**: More biologically motivated ViT-Retina performs as well as ViT-Blur on CIFAR-10 and leads it on both ImageNet sets, outperforming ViT-Blank under every impoverishment condition. ImageNet’s larger image size allows a more granular falloff of resolution, and it is more biologically faithful.

## 5 Spatial-frequency tunings diverge under impoverishment

We use narrow-band noise centered at different spatial frequencies (SFs) to determine how the fine-tuned models make the classification decision. A larger drop at an SF band indicates that the model depends on it more.

### Full condition

At pre-training mask ratio of 0.65 (Fig. 3A) all models share the same tuning *shape*, a peak at low-to-mid SF falling at high SF, but they differ in magnitude, with ViT-Blank sitting above every ViT-Blur. The ViT-Blur models are therefore less sensitive to noise, which agrees with noise robustness results (Fig. 2D). At pre-training mask ratio of 0.95 (Fig. 3B) the shapes themselves diverge: ViT-Blank’s peak shifts toward higher SFs and the ViT-Blur (strong) models shift to lower SFs. ViT-Blur (mild) shifts the least of all, maintaining the same shape as at pre-training mask ratio of 0.65. For ViT-Retina (Fig. 3C) the tuning shifts toward higher SF as the FOV widens: high-SF sensitivity rises at FOV 80° and 160°, and these are the two poorest performing ViT-Retina models (90.7% and 89.7% against 93.7% at FOV 5°).

### Short budget and reduced data

The same divergence appears under the other impoverishment conditions. Under the short budget (Fig. 3D) the SF tuning curve for ViT-Blank shifts to high SFs while the one for ViT-Blur holds its low-to-mid SF peak. Under reduced data (Fig. 3E) both curves shift toward higher SF peak, but the peak magnitude for ViT-Blank is much higher, so the models separate in high-SF magnitude rather than in peak location.

### The SF shift tracks the classification gap

Across every impoverishment condition, the model with lower classification accuracy is the one whose tuning has shifted further toward higher SF: the higher a model’s tuning centroid, the lower its downstream classification accuracy, with Pearson *r* = −0.73 across all 224 pre-trained models (Fig. 3F). Since the pre-training corruption type is the only thing that differs between the models, the observed link must be inherited from the pre-trained encoder, which we test next.

## 6 A closed-form linear theory of masked-reconstruction pre-training

### 6.1 The optimal linear encoder

Following Bisulco et al. (2025) and the classical linear-autoencoder analyses of Bourlard & Kamp (1988) and Baldi & Hornik (1989), we solve the *L*_2_ masked-reconstruction objective for a linear encoder of the corrupted input under ridge regularization. The derivation is given in full in App. C; it yields 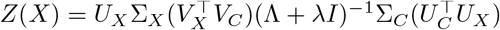, the representation the pre-trained encoder produces on a clean image, where 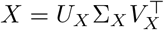 and 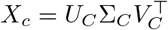 are the SVDs of the clean and corrupted training image matrices, 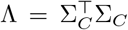, and *λ* is the ridge parameter. Every effect of corruption type, mask ratio, budget and training-set size must act through one of these factors. The key term is 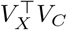, which projects the clean right singular basis onto the corrupted one: the impoverishment condition reshapes *V*_*C*_ and thereby selects which SF modes of *V*_*X*_ survive.

### 6.2 The linear encoder reproduces

#### ViT representation geometry and SF tuning

We first compare representational similarity matrices (RSMs) (Kriegeskorte et al., 2008) between the derived linear encoder and the pre-trained ViT on held-out CIFAR-10 (Figs. S27–S33 and App. D for detail). Each heat map is the pairwise cosine similarity between the encoder’s representations of every pair of corrupted images in the set, here 10 classes with two images each and 200 random corruption arrangements per image. Except for Linear-Blank and ViT-Blank at higher mask ratio, both the Linear and the ViT encoders show block-diagonal structure, so that differently corrupted versions of a given image map to mutually similar representations. The linear encoder predicts the empirical ViT RSM with a pooled Spearman correlation of 0.85 ± 0.01.

The encoder also reproduces every empirical pattern of §5 (Fig. 4 and App. D for detail): across all pre-training mask ratios and every impoverishment condition, Linear-Blur and Linear-Retina maintain lower SF reliance than Linear-Blank does. Pooled across all 224 pre-trained models and 14 SF centroids, the linear encoder predicts the ViT’s SF tuning at *r* = 0.88.

**Figure 4:**
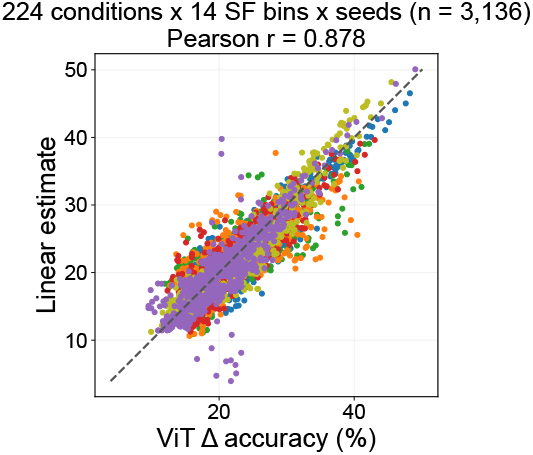
The linear theory predicts the ViT’s SF tuning. Linear-encoder SF tuning against the ViT’s accuracy drop.

### 6.3 Mechanism: singular-vector SF profile of the corrupted input

Having established that the linear theory approximates the ViT both in RSMs (Figs. S27–S33) and SF tuning (Fig. 4), we now localize the difference between corruption types using the derived linear encoder of §6.1. We do this in two stages: how SF content and class-discriminative signal are distributed across the modes of *V*_*X*_, and how much of each mode survives the encoder. Figure S34A plots the radially averaged power spectrum of each column of *V*_*X*_, with colour showing the singular value *σ*_*X*_. High *σ*_*X*_ columns concentrate at low SF and smaller *σ*_*X*_ columns carry progressively higher-SF content, so the basis is SF-ordered. The class-discriminative signal sits in the higher *σ*_*X*_ columns: the mean *d*^′^ across all 45 pairs of CIFAR-10 classes is well above the shuffled-label control there and converges to it for low *σ*_*X*_ (Fig. S34B). The high singular-value, low-SF modes of *V*_*X*_ therefore carry more class signal than the low singular-value, high-SF ones.

Figure S34C–F plots how much of each mode survives the encoder: 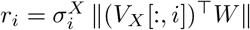, the magnitude of mode *i* against the SF centroid of *V*_*X*_ [:, *i*]. In the full condition at mask ratio of 0.65 (Fig. S34C), the magnitude across the SF centroid is similar among the different corruption types, consistent with their similar classification accuracy. At mask ratio of 0.95 (Fig. S34D), however, blank masking keeps the higher-SF modes while blur masking emphasizes the lower-SF modes that carry the class signal. Similarly, the models that transfer worst are the ones that keep the most high-SF modes (Fig. S34E and F, Figs. S35–S36). This is the mechanistic link from the pre-training corruption to the high-SF drift we measured in Fig. 3 and, ultimately, to the classification gap in Fig. 2.

## 7 Discussion

In this work, we experimented with replacing the random blank mask of masked-image modeling with band-limited ones: uniform blur at several Gaussian widths and cutoff frequencies, and retinal masking at several FOVs. At matched retained information, ViT-Blur and -Retina reconstruct fine details better than ViT-Blank, and their learned representations transfer better to downstream classification tasks. ViT-Blur and ViT-Retina are also more robust than ViT-Blank to impoverishment conditions, such as increased pre-trained mask ratio, limited pre-training epochs, limited training samples, and test time corruption. The gap widens further under test time input noise and common corruptions. The more biologically motivated ViT-Retina matches ViT-Blur on CIFAR-10 and outperforms it on ImageNet in downstream task. Noise probes and a closed-form linear encoder each point to the same cause: ViT-Blank relies more on high-SF information as the pre-training is more impoverished, while the biologically inspired ViT-Blur and ViT-Retina yield more robust visual representations that encode the class-information-carrying, high singular-value low-SF modes.

### Limitations

In the primate retina, cone density falls and rod density rises with increasing eccentricity, so along with resolution, color sensitivity also decays away from the fovea (Curcio et al., 1990). We did not implement this chromatic decay, and how it would affect representation learning is an open question. Second, ViT-Retina’s fixation is drawn uniformly at random, whereas humans sample a scene through a sequence of fixations that are goal directed (Najemnik & Geisler, 2005; Adeli et al., 2017). How this active sampling process changes the learned representation is worth testing. Third, we test our methods only on ViTs in this work, and we leave testing the proposed corruption techniques on other architectures such as CNNs (Pramod et al., 2022; Deza & Konkle, 2020; Jang & Tong, 2024) to future work.

## Reproducibility statement

All architecture and training hyperparameters are given in §3.2 and Appendix A. The corruption definitions, including the foveation falloff, are specified in §3.1, and the degrees-of-freedom accounting that matches the corruption types on retained information is given in App. A.3. The closed-form linear encoder is derived in full in Appendix C, and how its ridge parameter is selected, together with the leave-one-out cross-validation of that choice, is described in App. D. All evaluation probes are defined in §3.3 and use held-out data only. Code to reproduce the pre-training sweep, the probes and the linear theory will be released upon publication.

## Ethics statement

This work uses only standard public image benchmarks (CIFAR-10, ImageNet-100, ImageNet-1k) and involves no human subjects, no personally identifying data, and no deployed system. We note that ImageNet is known to carry annotation and representational biases, which any model pre-trained on it inherits.

## AI use statement

We used generative AI tools for language editing of author-written prose and for assistance with LaTeX formatting and figure-caption drafting. We did not use generative AI tools for research ideation, experimental design, data analysis, or the derivation in Appendix C. All experiments, results and numerical claims were produced and verified by the authors. We have reviewed all AI-assisted text and take responsibility for the final content of this work.

## Acknowledgments

Supported by Columbia University DSI Seed fund #202 and NIH/NEI grant R01 EY032938.

## A Additional method details

### A.1 Architecture and optimization

Table 2 summarizes the architecture and the key training hyperparameters. Fine-tuning uses the standard recipe (AdamW, RandAugment, Mixup/CutMix, label smoothing 0.1) applied only to clean images, and is identical across corruption types so that any difference in downstream accuracy is attributable to pre-training alone. Both ImageNet configurations are the standard SimMIM ViT-B/16 recipe with no modification other than the corruption type, the mask ratio and the number of pre-training epochs.

**Table 2:** Architecture and training hyperparameters. Both ImageNet columns are the unmodified SimMIM ViT-B/16 configuration, at 800 pre-training epochs on ImageNet-100 and 100 on ImageNet-1k. ImageNet learning rates follow the linear scaling rule, LR = base × (global batch)*/*512, giving 2.5 × 10^−5^ (pre-training) and 3.125 × 10^−4^ (fine-tuning) at global batch 128. Fine-tuning uses drop-path 0.1 and gradient clipping at 5.0, and the reconstruction loss is taken over the whole image for every corruption type. ^†^The retinal-masking models on either ImageNet set use 16 × 16 mask patches; every other corruption type uses 32 × 32.

|  | CIFAR-10 | ImageNet-100 | ImageNet-1k |
| --- | --- | --- | --- |
| Image size | $32 \times 32$ | $224 \times 224$ | $224 \times 224$ |
| Patch size (model) | $4 \times 4$ | $16 \times 16$ | $16 \times 16$ |
| Patch size (mask) | $4 \times 4$ | $32 \times 32^{\dagger}$ | $32 \times 32^{\dagger}$ |
| Embed dim | 384 | 768 | 768 |
| Depth / heads | 6 / 12 | 12 / 12 | 12 / 12 |
| Pre-train epochs | 200 (warmup 7) | 800 (warmup 10) | 100 (warmup 10) |
| Fine-tune epochs | 100 (warmup 7) | 100 (warmup 20) | 100 (warmup 20) |
| Optimizer | AdamW | AdamW | AdamW |
| Base LR (pre / ft) | $2 \times 10^{-3} / 2 \times 10^{-2}$ | $1 \times 10^{-4} / 1.25 \times 10^{-3}$ | $1 \times 10^{-4} / 1.25 \times 10^{-3}$ |
| Weight decay | 0.08 | 0.05 | 0.05 |
| LR schedule (pre / ft) | multi-step / cosine | multi-step / cosine | multi-step / cosine |
| Layer decay (ft) | — | 0.65 | 0.65 |
| Batch size | 64 | 128 | 128 |

**Table 3:**
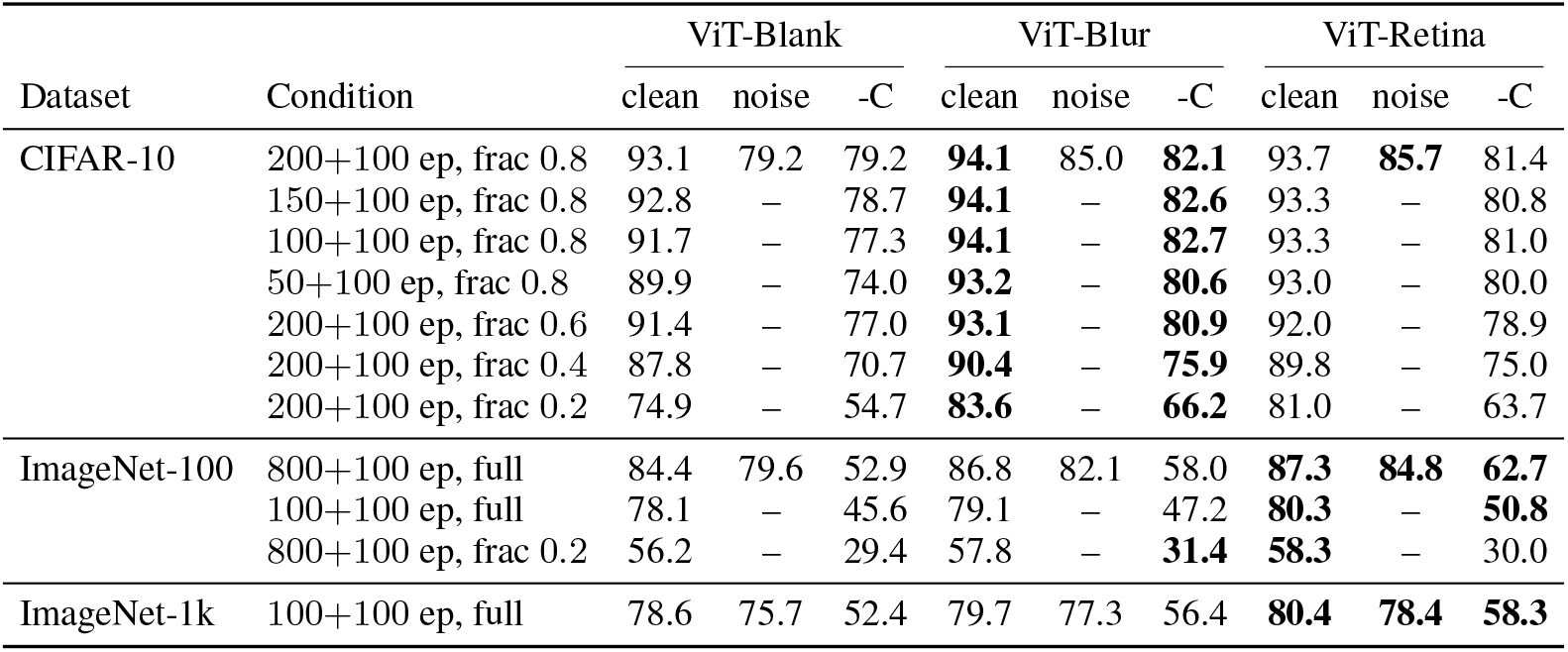
Clean, noise and corruption accuracy for every impoverishment condition. Conditions are given as pre-training+fine-tuning epochs and training fraction. Each entry is the best-clean-accuracy setting of that corruption type, reported under three evaluations. *clean*: accuracy on the clean test set. *noise*: accuracy when Gaussian pixel noise (*µ* = 0, *σ* = 0.05 on [0, 1] pixel values) is added to the test image at evaluation only, measured in the full conditions only. *-C*: mean accuracy on the matching corruption benchmark (CIFAR-10-C, ImageNet-100-C or ImageNet-1k-C), averaged over 15 corruptions at 5 severities (Hendrycks & Dietterich, 2019). Neither noise nor corruption is ever applied during pre-training or fine-tuning. Best in each column triple in bold.

### A.2 Retinal masking

Each retinal-masking sample designates one patch as foveal, drawn uniformly at random over the image and snapped to a patch centre, with a new fixation drawn on every forward pass. Every patch is then low-pass filtered at the cutoff *f*_*c*_(*e*) = *f*_0_*/*(1 + *e/e*_*h*_), evaluated at the eccentricity *e* of that patch’s centre from the fixation, with Gaussian width *σ*_*f*_ = 0.5 *f*_*c*_.

Here *e*_*h*_ = 2.3° is the eccentricity at which human resolution falls to half its foveal value (Geisler & Perry, 1998), and setting *f*_0_ to the Nyquist frequency leaves the foveal patch unfiltered. A retinal-masked input is therefore a single glimpse that is sharp at one point and progressively blurred away from it.

The field of view (FOV) converts pixels to degrees: *e* is scaled so that the diagonal corners of the image subtend the FOV. A larger FOV means more degrees per pixel, so the cutoff falls faster in pixel terms and the input is *more* severely blurred, with FOV 160° the harshest setting and FOV 2.5° the mildest. The cutoff is floored at a minimum value (1 cyc/image) and it is quantized to integer cycles per image during training. Figure S1 plots the resulting profile for every FOV.

**Figure S1:**
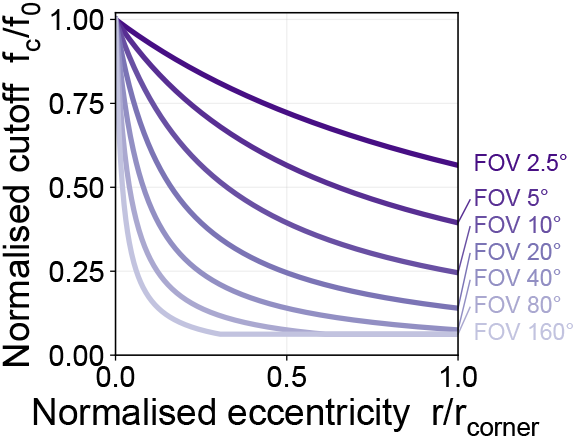
The eccentricity-dependent cutoff of the retinal-masking models.

### A.3 Degrees-of-freedom accounting

Retained DOF has two parts: the pixels that survive at full resolution, and the Fourier coefficients that survive the low-pass filter.

The first part counts the pixels of the visible, uncorrupted patches. A visible patch contributes its own area, so 16 pixels for CIFAR-10’s 4×4 mask patches and 1024 for the 32×32 mask patches used on ImageNet, or 256 for the retinal-masking models on either ImageNet set, which use 16×16.

The second part counts the Fourier coefficients the filter leaves non-zero. A cutoff of *c* cyc/image zeroes every coefficient whose radial frequency is *c* cyc/image or higher, so the surviving coefficients are the lattice points (*u, v*) with 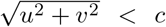. At cutoff 2 these are {(0, 0), (0, ±1), (±1, 0), (±1, ±1)}, nine points in all. Counting this way, a cutoff of 2 cyc/image leaves 9 coefficients, a cutoff of 3 leaves 25, and mild blur, whose cutoff is 8 cyc/image, leaves 193.

The two parts add. On CIFAR-10, a cutoff-2 blur with three visible patches therefore retains 9 + 16 × 3 = 57 degrees of freedom (an upper bound, since the low frequencies are properties of the whole image and it counts the visible patches a second time), against 48 for blank masking with the same three visible patches, which passes no low frequencies and so contributes visible pixels only.

For the retinal-masking models the passband count is area-weighted over patches, since each patch has its own cutoff, and averaged over fixations, with the standard deviation over fixations shown as a horizontal error bar in Fig. S5.

### A.4 Evaluation probes

The *SF band-limited noise probe* constructs noise by multiplying white noise in the Fourier domain by a Gaussian ring of standard deviation 0.52 cyc/image centred at the target frequency, scaled to *σ* = 0.06 on [0, 1] pixel values. Noise is added to 1,000 validation images at 14 ring centres from 1 to 15 cyc/image, with 10 independent noise draws per centre; we report the drop in top-1 accuracy relative to the clean images. The tuning *centroid* used in Fig. 3F is the drop-weighted mean ring centre, 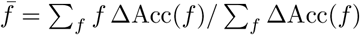.

The *corruption probes* use the standard 15 corruption types at severities 1–5: CIFAR-10-C over the 10,000 CIFAR-10 test images per corruption and severity, ImageNet-100-C over 5,000 validation images, and ImageNet-1k-C over 30,000 images. Reconstruction error is masked-region *L*_1_ in normalized pixel units, averaged over 30 random mask (and fixation) draws on a fixed held-out image set.

## B Compute resources

All models were trained on NVIDIA L40S GPUs, with memory and storage from an internal cluster.

## C Derivation of the closed-form linear encoder

A linear encoder is a useful proxy for the ViT because the two agree to first order. Let *δX* be the band-limited noise added by the SF probe (§3.3). A first-order Taylor series expansion of the trained ViT encoder *f* gives *f* (*X* + *δX*) − *f* (*X*) ≈ *δX J* (*X*), where the Jacobian *J* (*X*) is itself a linear transform, while a linear encoder *Z*(*X*) = *XW* satisfies *Z*(*X*+*δX*) − *Z*(*X*) = *δX W*. Heuristically, if the two encoders produce the same representation for both the image and its perturbation, the two differences are equal, so *δX J* (*X*) ≈ *δX W* for arbitrary *δX* and hence *J* (*X*) ≈ *W*. The linear encoder may therefore approximate the ViT’s response to the probe.

### Setup

Let there be *M* clean training images, each corrupted *K* times with independent mask (or fixation) realizations. Denote by **x**_*c,i,j*_ ∈ ℝ^*n*^ the *j*-th corrupted version of the *i*-th image and stack all corrupted inputs row-wise into *X*_*c*_ ∈ ℝ^*MK*×*n*^. Let *X* ∈ ℝ^*MK*×*n*^ be the matrix whose ((*i*−1)*K* + *j*)-th row is the clean image **x**_*i*_, so each clean image appears *K* times aligned with its *K* corrupted copies; this alignment is what makes ∥*X* − *ZD*∥_*F*_ a per-sample reconstruction loss. Let *Z* ∈ ℝ^*MK*×*d*^ be the encoder’s last-layer representation of *X*_*c*_ and *D* ∈ ℝ^*d*×*n*^ the linear decoder. We use the *L*_2_ masked-reconstruction objective

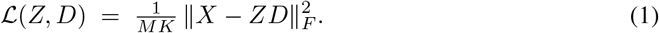

### Step 1: optimal decoder for fixed *Z*

Expanding via 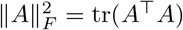,

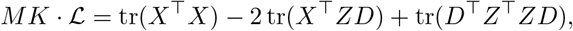

using tr(*A*) = tr(*A*^⊤^) to combine the cross terms. With *∂* tr(*A*^⊤^*B*)*/∂B* = *A* and *∂* tr(*B*^⊤^*AB*)*/∂B* = (*A* + *A*^⊤^)*B* (here *A* = *Z*^⊤^*Z* is symmetric), 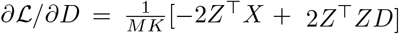. Setting this to zero gives the normal equation *Z*^⊤^*ZD* = *Z*^⊤^*X*, i.e.

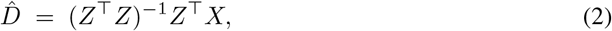

assuming *Z*^⊤^*Z* invertible (generically true once *MK* ≥ *d* and *Z* has full column rank).

### Step 2: reduced objective in *Z*

Substituting equation 2 into equation 1 and writing *P*_*Z*_ := *Z*(*Z*^⊤^*Z*)^−1^*Z*^⊤^ for the orthogonal projector onto the column space of *Z*, we have 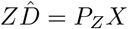 and hence 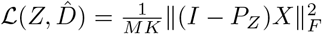. Using 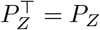 and 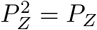.

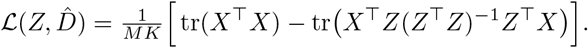

The first term does not depend on *Z*, so minimizing ℒ is equivalent to

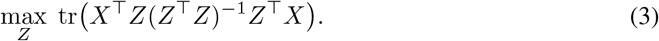

### Step 3: optimal 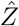 under orthonormality

Objective equation 3 is invariant under *Z* ↦ *ZA* for any invertible *A*, since *ZA*(*A*^⊤^*Z*^⊤^*ZA*)^−1^*A*^⊤^*Z*^⊤^ = *P*_*Z*_; it depends only on colspan(*Z*). To select a representative we impose *Z*^⊤^*Z* = *I*_*d*_, under which the objective reduces to tr(*Z*^⊤^*XX*^⊤^*Z*). Forming the Lagrangian *J* = tr(*Z*^⊤^*XX*^⊤^*Z*) − tr(Ω(*Z*^⊤^*Z* − *I*_*d*_)) with symmetric multipliers Ω and setting *∂J /∂Z* = 0 gives *XX*^⊤^*Z* = *Z*Ω; left-multiplying by *Z*^⊤^ yields Ω = *Z*^⊤^*XX*^⊤^*Z*. Substituting the 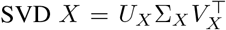, so that 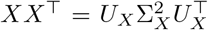, the choice 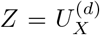 (the first *d* columns of *U*_*X*_) gives 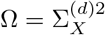, which is diagonal and hence symmetric as required. This is a valid stationary point and we take it as the canonical maximizer,

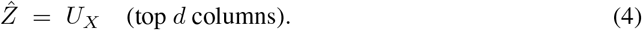

### Step 4: linear encoder and ridge regularization

We now replace the implicitly defined *Z* with a *linear encoder* of the corrupted input, *Z* = *X*_*c*_*W*, and seek the *W* whose induced *Z* best approximates 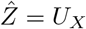. Training uses weight decay, which is exactly a ridge penalty, so

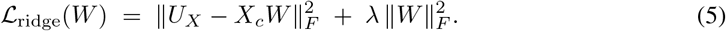

Expanding and using 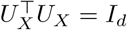,

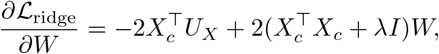

so the regularized normal equation gives

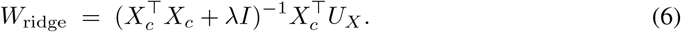

The matrix 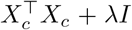 is positive definite for any *λ >* 0, so the inverse always exists — one practical reason for regularizing. As *λ* → 0 this reduces to the unregularized least-squares solution where that exists.

### Step 5: spectral form

Take the full SVD 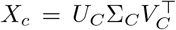, so that 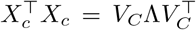 with 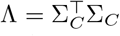 and *V*_*C*_ orthogonal. Then 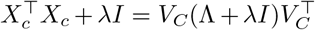 and 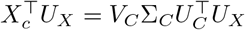. Substituting into equation 6 and using 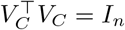,

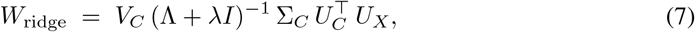

which is the closed-form solution quoted in §6.1.

### Step 6: induced representation on a clean input

Applying equation 7 to a clean input matrix 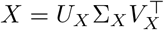,

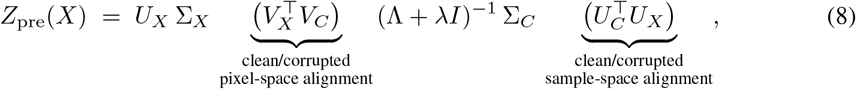

which is the induced representation *Z*(*X*) quoted in §6.1: the closed-form prediction for the representation the encoder produces on a clean image after masked-reconstruction pre-training. Every effect of corruption type, mask ratio, epoch budget and training-set size must act through one of these factors, and the analyses of §6.3 proceed by asking which.

## D Linear theory: extended results

This appendix gives the representational similarity matrices, the per-model tuning fits, and the control analyses behind Figs. 4 and S34 of the main text.

### D.1 Representation geometry

The linear encoder is useful as a theory of the ViT only if the pre-training representation it predicts is geometrically similar to that of the trained ViT. We test this by comparing the representational similarity matrices (RSMs) (Kriegeskorte et al., 2008) of *X*_*c*_*W* on held-out CIFAR-10. We computed RSMs across many held-out image pairs and corruption types and observed the same trend throughout; the matrices shown use 10 classes with two images each and 200 random corruption arrangements per image (Figs. S27–S33). Each RSM is the pairwise cosine similarity between the encoder’s representations of every pair of corrupted images in the set. The optimal encoder 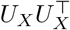 gives a block-diagonal structure, in that the 200 differently corrupted versions of a *given* image map to mutually similar representations, forming one block per image. At lower mask ratio both the linear ridge readout and the empirical ViT reproduce this structure under every corruption type. At higher mask ratio the block structure breaks down in both Linear-Blank and ViT-Blank, so that the encoder can no longer tell corrupted versions of one image from another, while blur and retinal masking keep it intact in both. Pooled over all 224 pre-trained models the Linear–ViT Spearman correlation is 0.85 ± 0.01, with the per-model value given in each panel title. The linear ridge encoder is therefore a reasonable surrogate for the ViT at the level of pre-training representation geometry, which licenses using it on clean inputs to localize the differences between corruption types in SF tuning.

### D.2 What the theory predicts

The derived latent representation contains the term 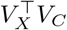, which projects the clean right singular basis onto the corrupted one in pixel space. The core prediction is that the impoverishment conditions reshape *V*_*C*_ and thereby select which spatial-frequency modes of *V*_*X*_ are preserved in the learned representation. *V*_*C*_ depends on corruption type, mask ratio, the number of mask arrangements per image, and the number of training samples per class, and is shaped differently for blank than for blur along each of these axes; Figs. S35–S36 show how *V*_*C*_ changes with each factor.

### D.3 SF tuning of the linear encoder

For a linear encoder *Z* = *XW* and the same band-limited probe *δx* as §3.3, the SF sensitivity at a given band is 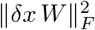. We select the ridge parameter *λ* per pre-trained model to maximize the Pearson correlation between the linear prediction and the empirical SF tuning, which fixes the shape of the tuning curve, and then apply a single scaling to match the accuracy-drop range. Across every impoverishment condition the linear encoder reproduces the empirical pattern. At high mask ratio the peak of Linear-Blank drifts to high SF while Linear-Blur drifts to *lower* SF, matching Fig. 3A–B. Under a short budget Linear-Blank again drifts to high SF while Linear-Blur holds its low-to-mid peak. Under reduced data both drift to high SF, but Linear-Blank drifts much further. Per-model fits, with the fitted *λ*^∗^ and the per-model Pearson correlation in each title, are given in Figs. S20–S26; pooled across all 224 pre-trained models and 14 ring centres the linear theory predicts the empirical ViT SF tuning at *r* = 0.88 over 224 × 14 = 3,136 points (Fig. 4).

### D.4 The fitted *λ* is not free

Figure S37 addresses the concern that fitting *λ* per model could make the agreement with the ViT trivial. Across all 224 pre-trained models the fitted *λ*^∗^ scales monotonically with log tr(*X*_*c*_), following a smooth sigmoid (Spearman *ρ* = 0.84), so *λ* tracks a bulk property of the corrupted input rather than absorbing per-model idiosyncrasy. Under leave-one-out cross-validation, fitting the sigmoid to the other 223 models, predicting the held-out *λ* from its tr(*X*_*c*_) alone, and scaling the resulting curve, the linear encoder still predicts the ViT’s tuning at *r* = 0.61 over the same 3,136 points. The tuning is therefore predictable from the corrupted training set alone, with no free parameter fitted to the ViT.

**Figure S2:**
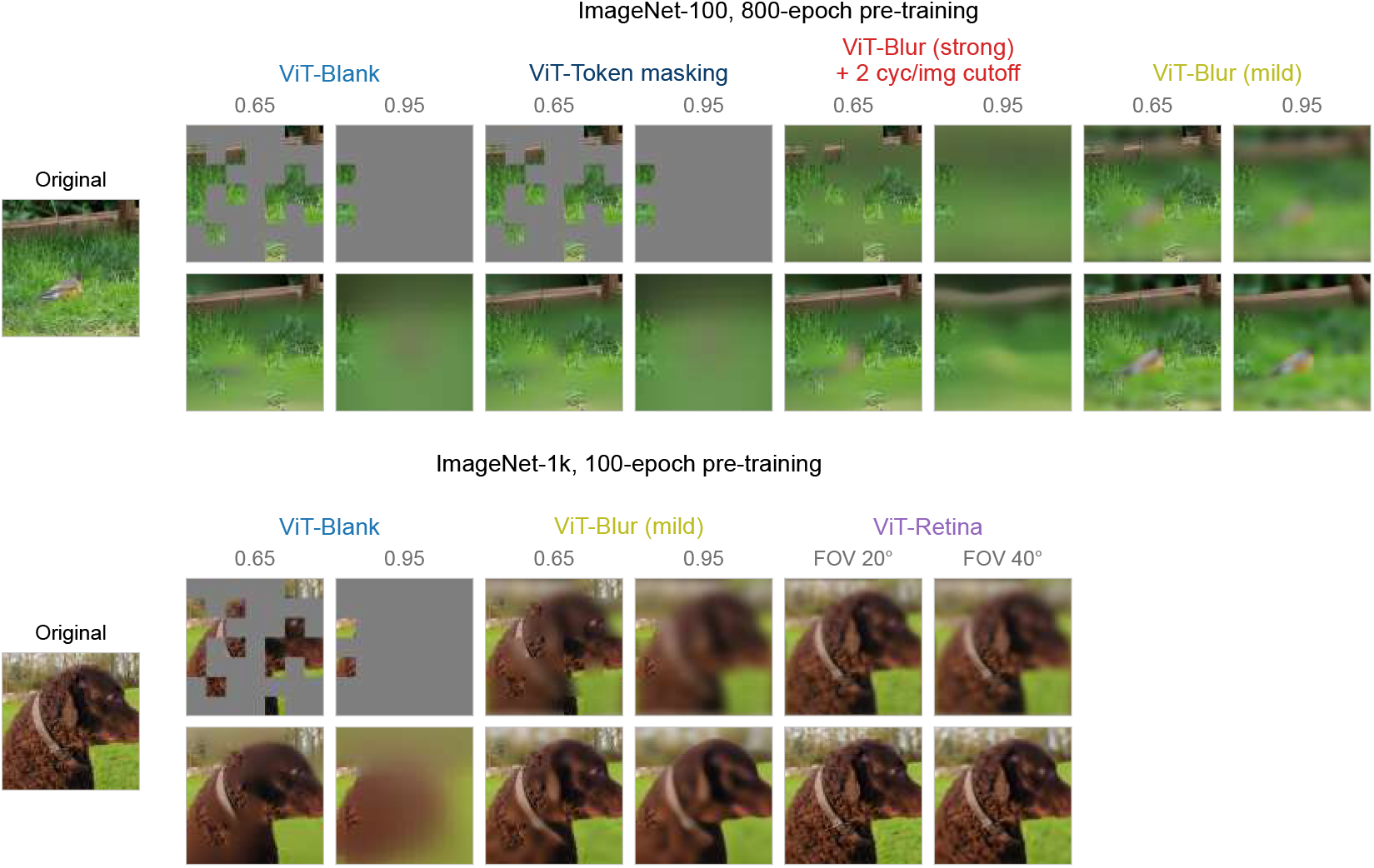
Inputs and reconstructions on ImageNet. Top: ImageNet-100 at 800 epochs, showing ViT-Blank, token masking, ViT-Blur (strong + 2 cyc/image) and ViT-Blur (mild) at mask ratios 0.65 and 0.95. Bottom: ImageNet-1k at 100 epochs, showing ViT-Blank and ViT-Blur (mild) at mask ratios 0.65 and 0.95, and ViT-Retina at FOV 20° and 40°. Originals at the left, corrupted inputs on the upper row of each block and each model’s reconstruction below.

**Figure S3:**
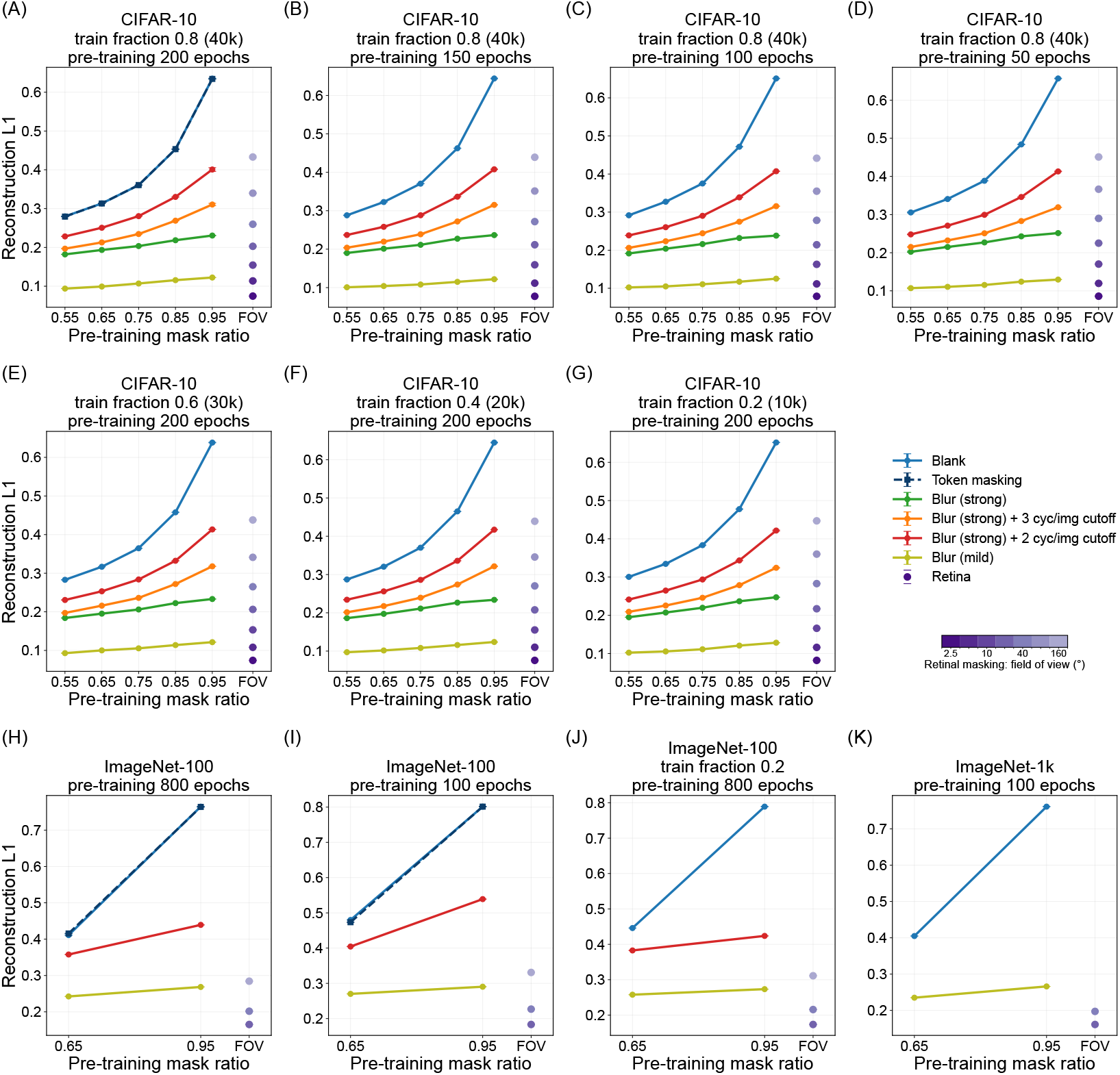
Reconstruction *L*_1_ against the pre-training mask ratio for every impoverishment condition. (CIFAR-10 budgets and fractions; ImageNet-100 at 800 epochs, 100 epochs and fraction 0.2; ImageNet-1k at 100 epochs). Mean ± SEM over 30 mask draws; ViT-Retina in the FOV column.

## E The role of the sharp fovea at matched information

This appendix gives the reconstruction figures referenced in §4. Figure S4 additionally asks whether a sharp region is needed at all, by bracketing blank masking in retained DOF with fully blurred inputs, blur plus a few sharp patches, and retinal masking, and scoring each on both reconstruction error and fine-tuned classification accuracy. The two orderings differ: the fully blurred inputs are reconstructed second best but transfer worst, while retinal masking leads on both at the lowest DOF of any model. What transfers is therefore a sharp fovea with a coarse surround, not ease of reconstruction or the amount of information retained.

**Figure S4:**
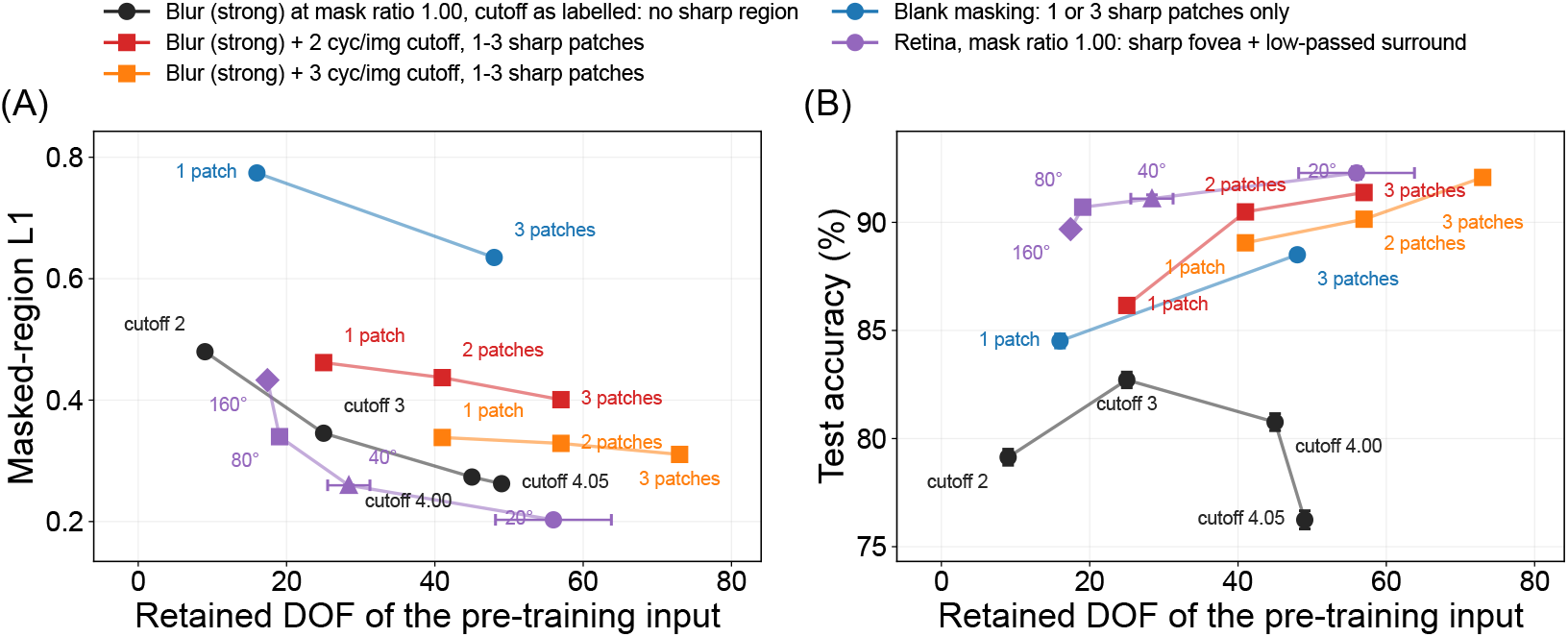
What a sharp region buys at matched information. Full-condition CIFAR-10 models against the retained DOF of their pre-training input, lines joining members of a family: uniform blur at mask ratio 1.0 (black; no sharp pixel anywhere), blur with cutoff 2 (red) or 3 (orange) plus 1–3 sharp patches, blank masking with 1 or 3 sharp patches and nothing else (blue), and ViT-Retina at FOV 160–20° (purple; one sharp fovea plus a graded surround). **(A)** Masked-region reconstruction *L*_1_ (mean ± SEM over 30 mask draws): the retinally masked inputs are reconstructed best, then the fully blurred ones, then blur with sharp patches, and blank worst. **(B)** Fine-tuned validation accuracy, in a different order: ViT-Retina best, then blur with sharp patches, then blank, with the fully blurred inputs worst. At equal patch count a low-passed surround is worth 1.7–4.6 points over blank with one patch and 2.9–3.6 with three; the retinally masked inputs are highest at the lowest DOF.

**Figure S5:**
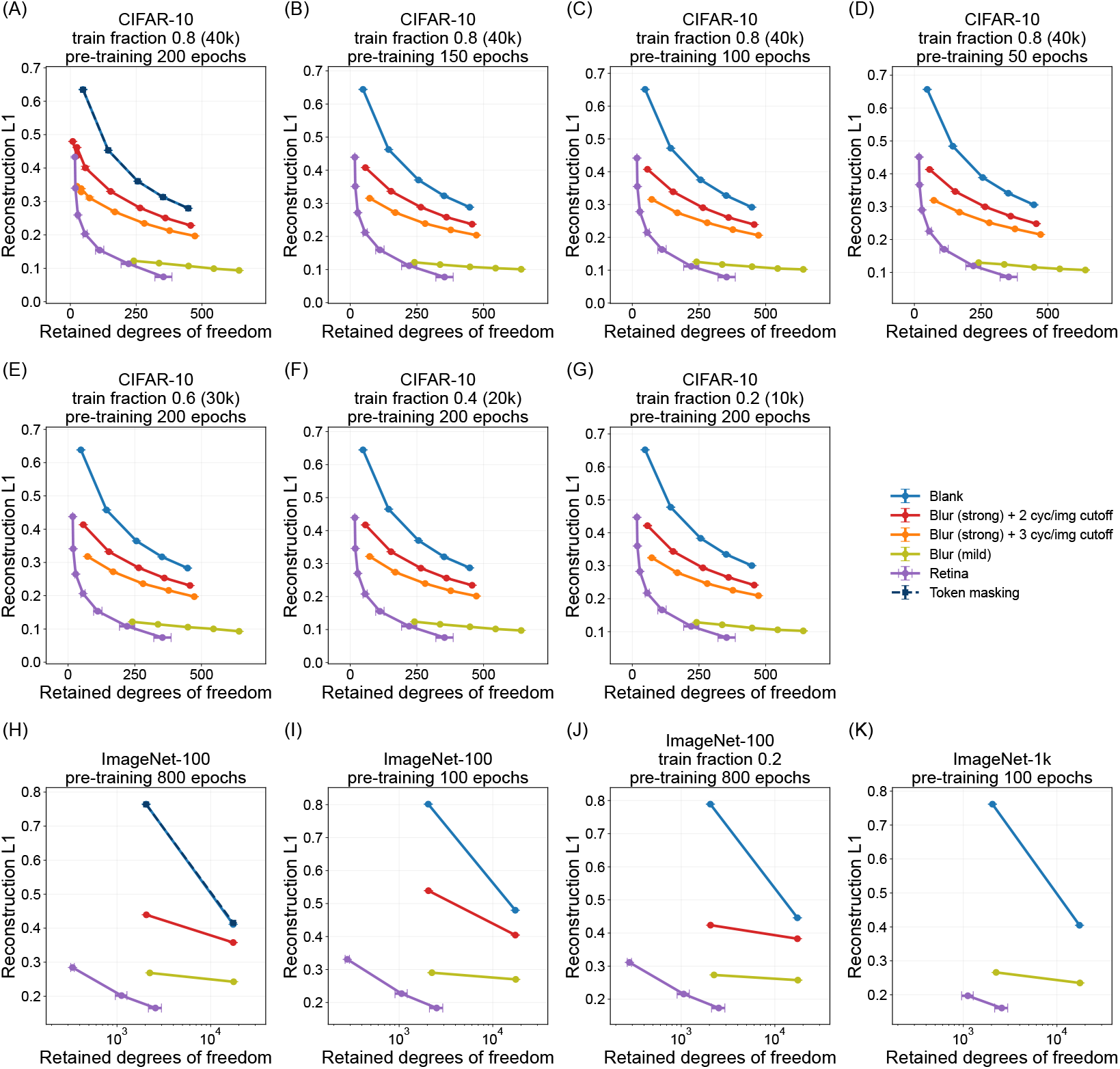
Reconstruction *L*_1_ against retained degrees of freedom for every impoverishment condition of Fig. S3. Along each masked curve the mask ratio falls from left to right, and along each retinal-masking curve the field of view narrows: from 0.95 to 0.55 and from 160° to 2.5° on CIFAR-10, and from 0.95 to 0.65 and from 160° to 20° on ImageNet-100. In the CIFAR-10 full condition (A), the ViT-Blur (strong) + 2 cyc/image and + 3 cyc/image cutoff curves extend further left, to mask ratio 1.0, passing through points with one visible patch and then no visible patch, so that they bracket blank masking at mask ratio 0.95. ImageNet-1k contributes two points per corruption type, at mask ratios 0.95 and 0.65 and at FOV 40° and 20°.

**Figure S6:**
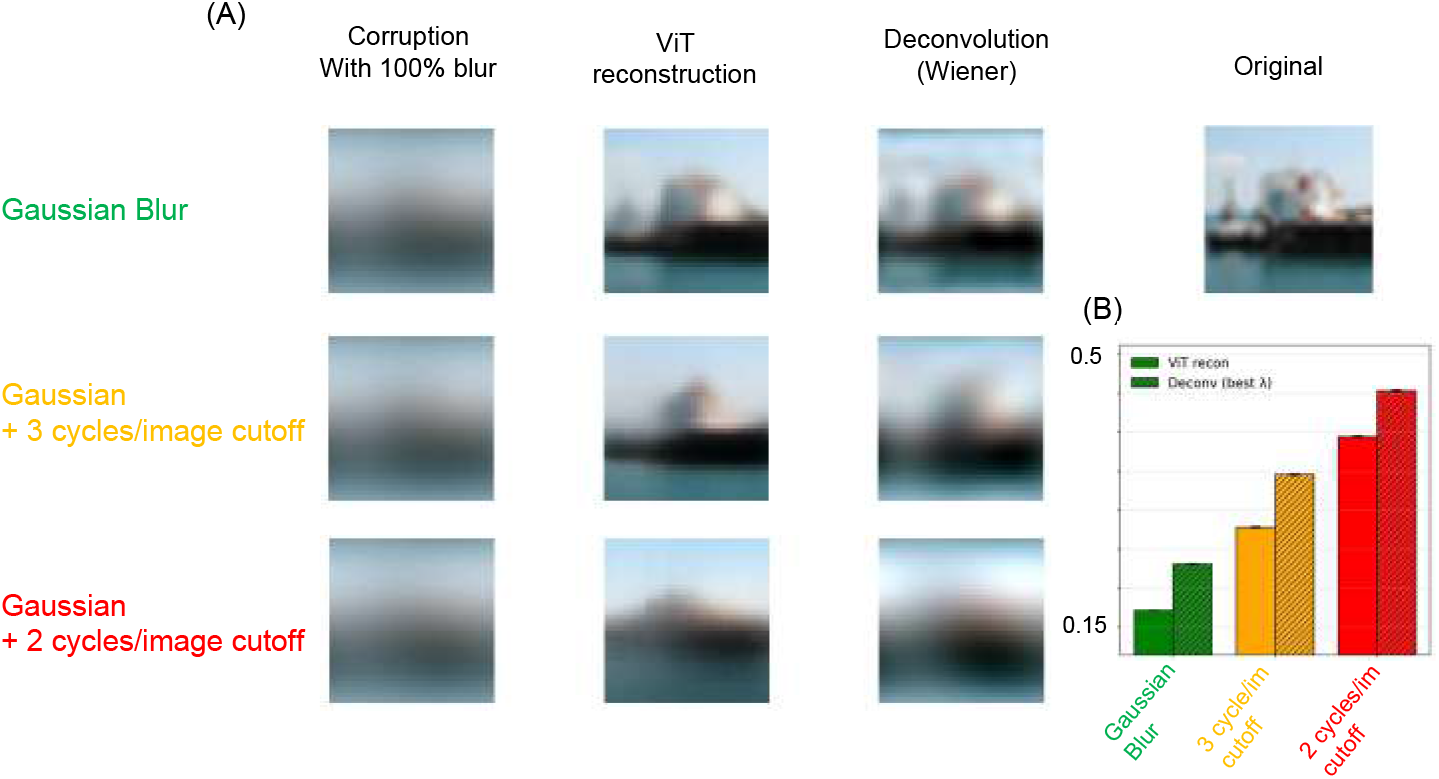
Reconstruction of a fully blurred image by the ViT compared with linear (Wiener) deconvolution. Evaluated at mask ratio 1.0, where every patch is blurred and no sharp patch is left to infer from. The Wiener filter (`skimage.restoration.wiener`) is given the ground-truth blur kernel and inverts it explicitly, whereas the ViT never sees the kernel. **(A)** Example CIFAR-10 images under each blur corruption type; columns, left to right: the corrupted input, the ViT reconstruction, and the Wiener deconvolution. **(B)** Reconstruction *L*_1_ error against the clean image for the two methods in each blur corruption type.

## F Classification and corruption robustness: extended results

Figure S7 reports the two-parameter blur sweep from which the main-text blur conditions were selected. Accuracy rises with *σ*_*f*_ for every cutoff, and mild blur (*σ*_*f*_ = 4, cutoff 8) is the best cell at 93.8%. The remaining figures of this appendix give the clean-accuracy, noise and corruption sweeps in full, on both the mask-ratio and the retained-DOF axis.

**Figure S7:**
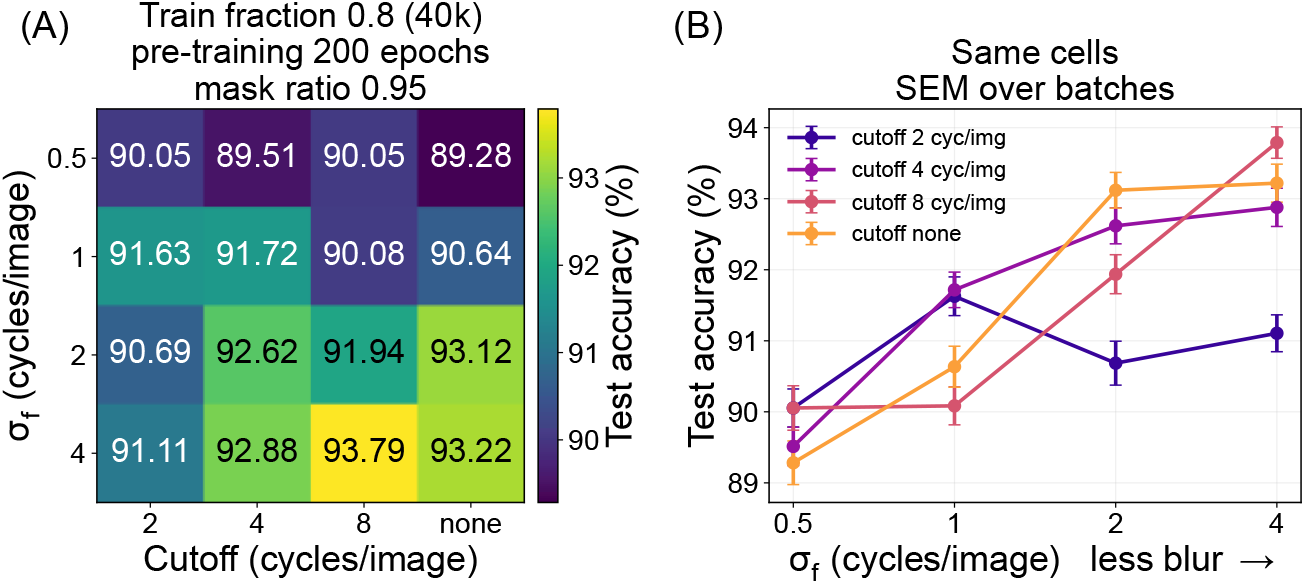
The blur parameter sweep at mask ratio 0.95, full condition. **(A)** Validation accuracy over *σ*_*f*_ (0.5–4 cyc/image) cutoff × (2, 4, 8, none). **(B)** The same cells as lines against *σ*_*f*_, with batch SEM. Less blur is better; *σ*_*f*_ = 4 with cutoff 8 is the best cell.

**Figure S8:**
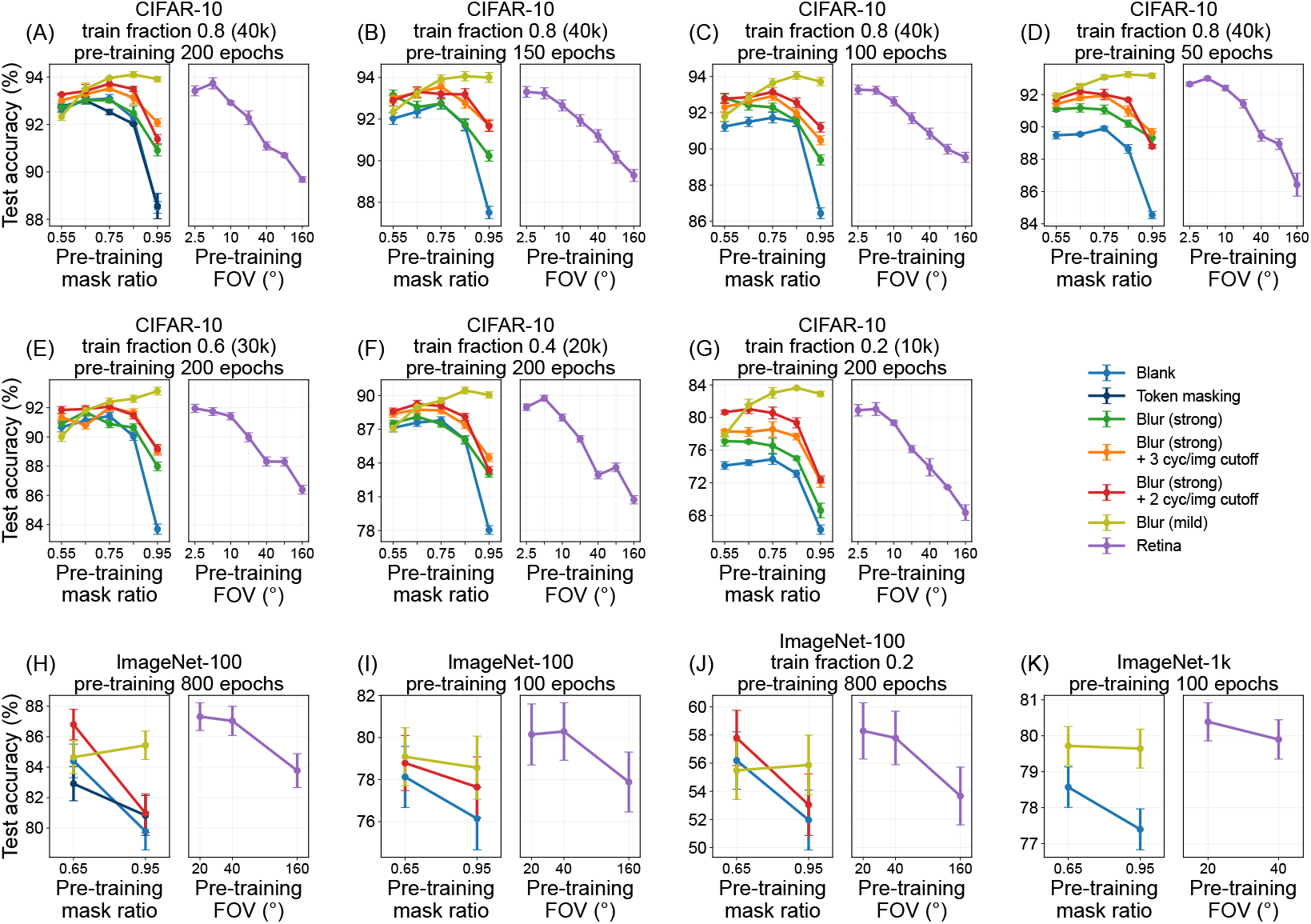
Clean accuracy for every condition across pre-trained mask ratio and FOV.

**Figure S9:**
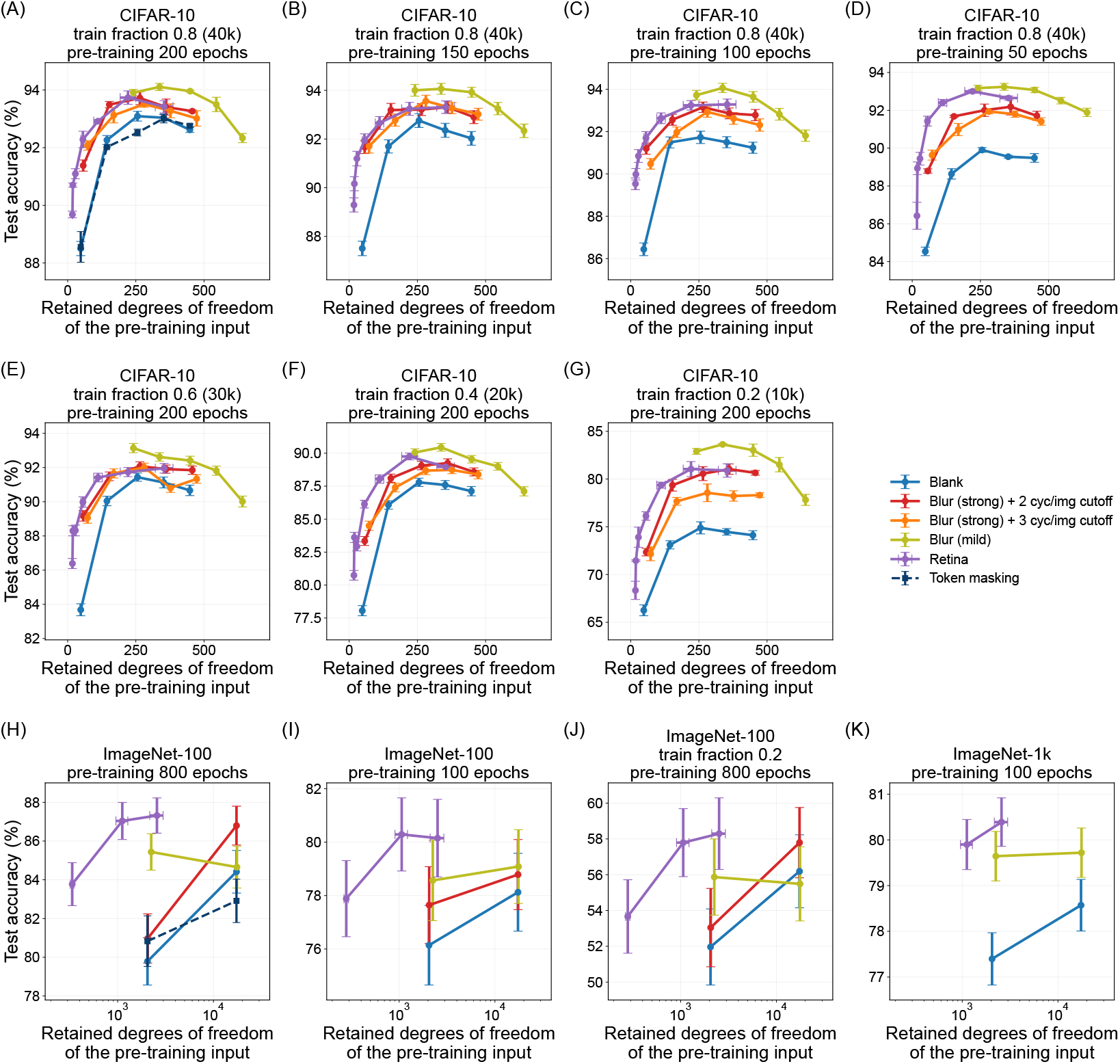
Fig. S8 against the retained degrees of freedom of the pre-training input. Along each curve the pre-training mask ratio and the field of view fall from left to right: from 0.95 to 0.55 and from 160° to 2.5° on CIFAR-10, and from 0.95 to 0.65 and from 160° to 20° on ImageNet-100. ImageNet-1k contributes two points per model, at mask ratios 0.95 and 0.65 and at FOV 40° and 20°.

**Figure S10:**
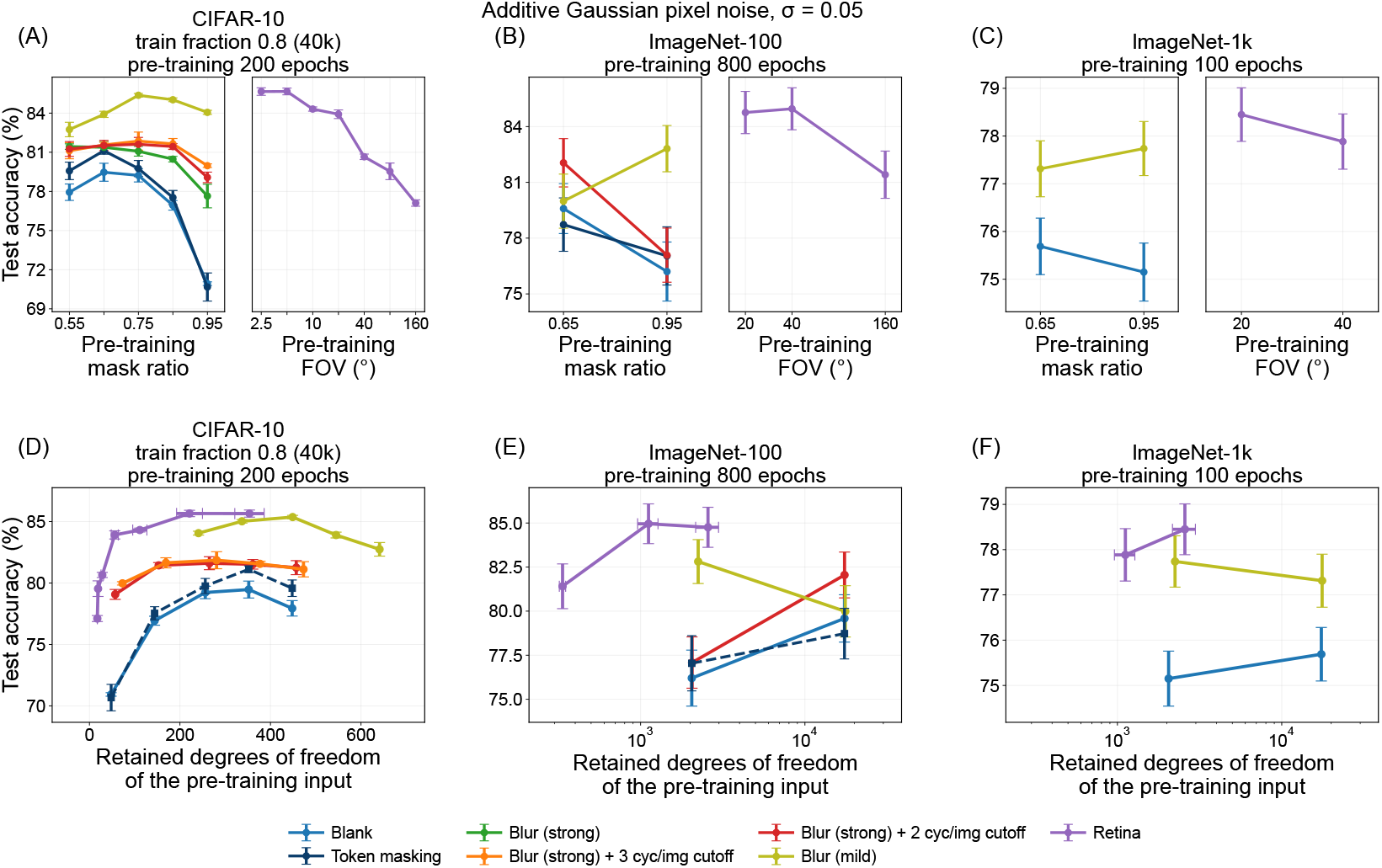
Accuracy under additive Gaussian pixel noise on every dataset, applied at evaluation only, for the models of the full condition, against the pre-training mask ratio and the field of view. Noise is never applied during pre-training or fine-tuning.

**Figure S11:**
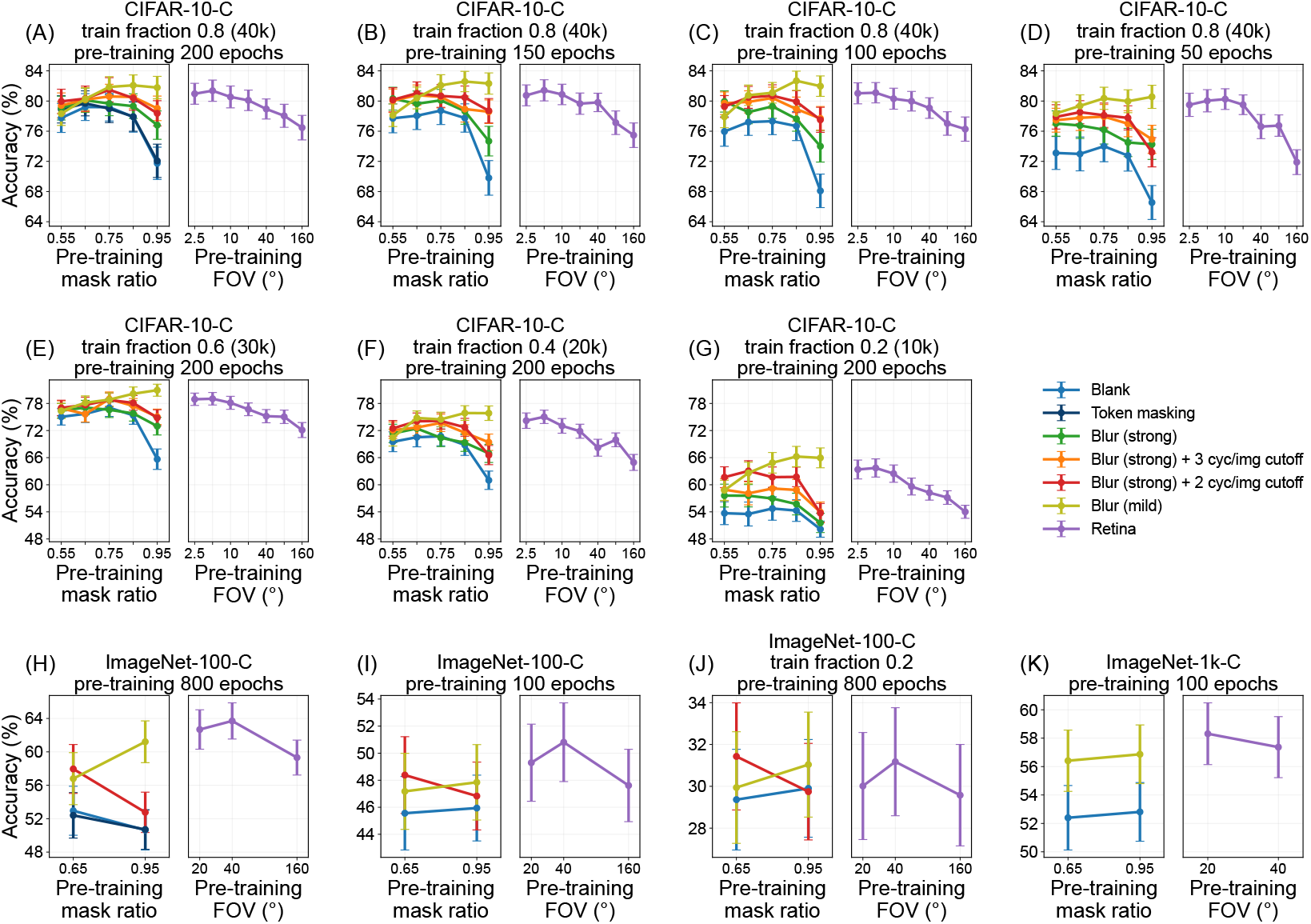
Corruption robustness on every dataset: mean accuracy over the 15 standard corruptions and 5 severities against the mask ratio and the field of view. Error bars: SEM across corruption types.

**Figure S12:**
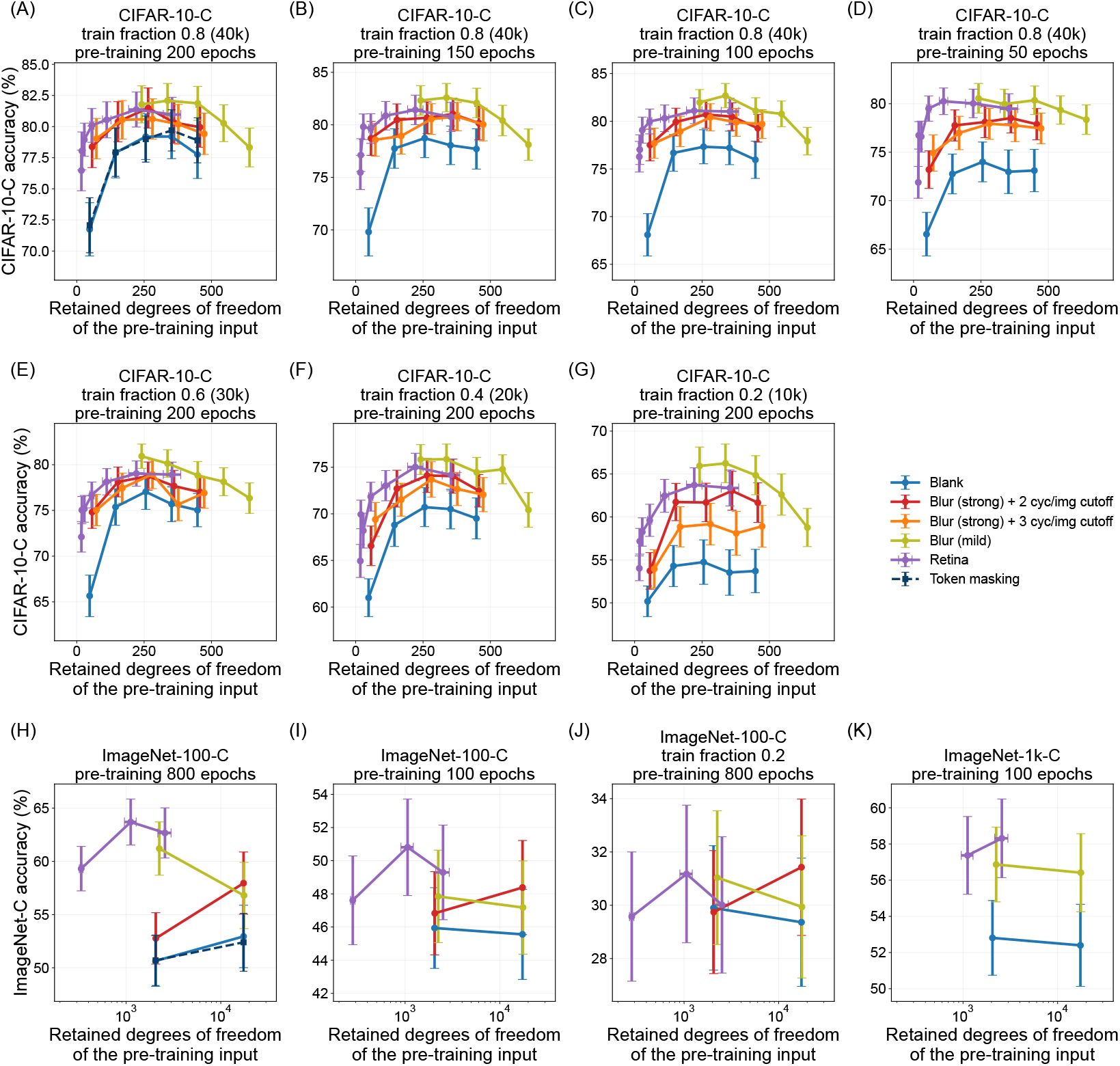
The panels of Fig. S11 with the retained degrees of freedom of the pre-training input on the *x*-axis.

**Figure S13:**
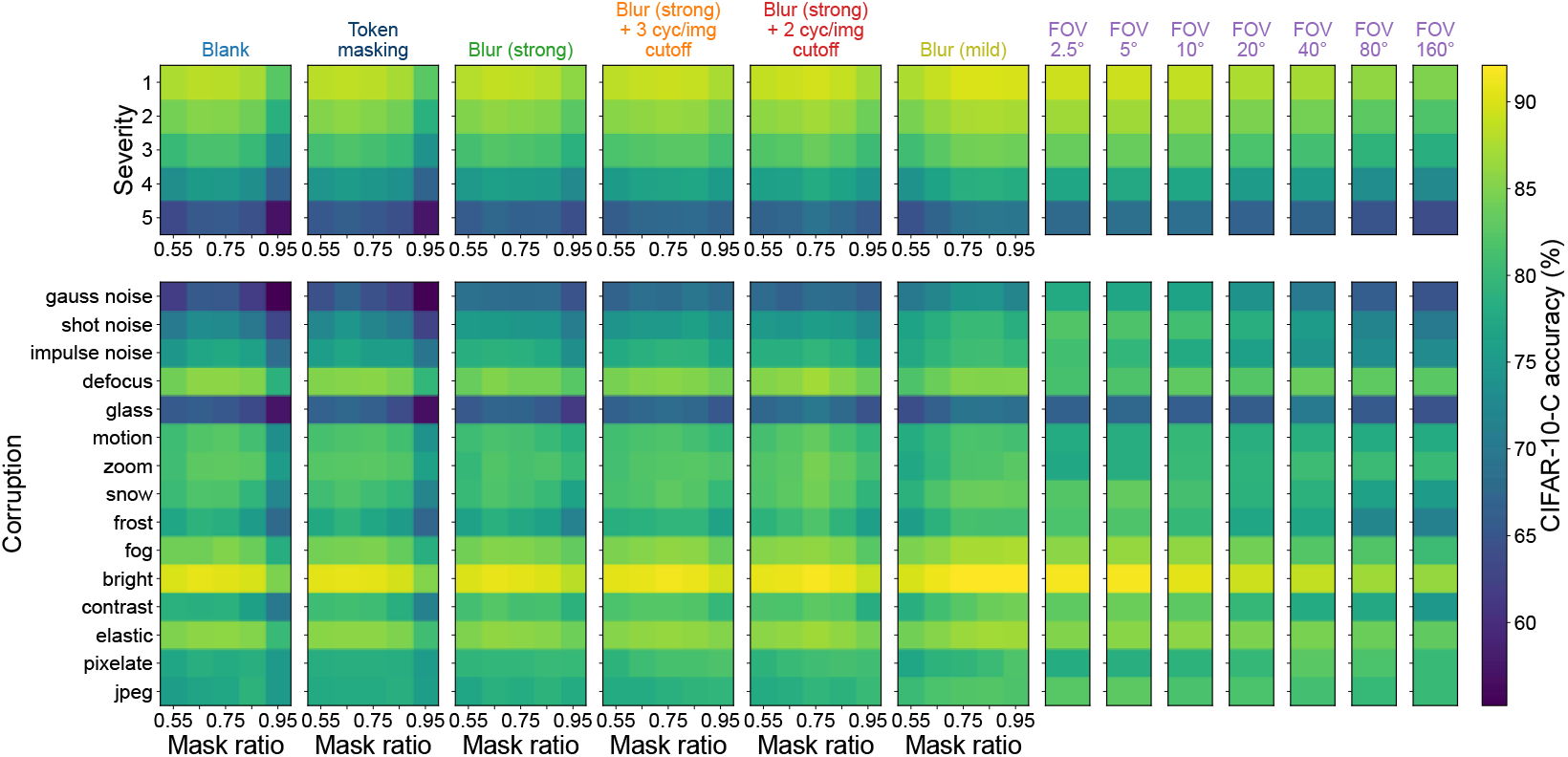
CIFAR-10-C by severity (top) and by corruption type (bottom) against the mask ratio, one column per arm, full condition; one shared colour scale.

**Figure S14:**
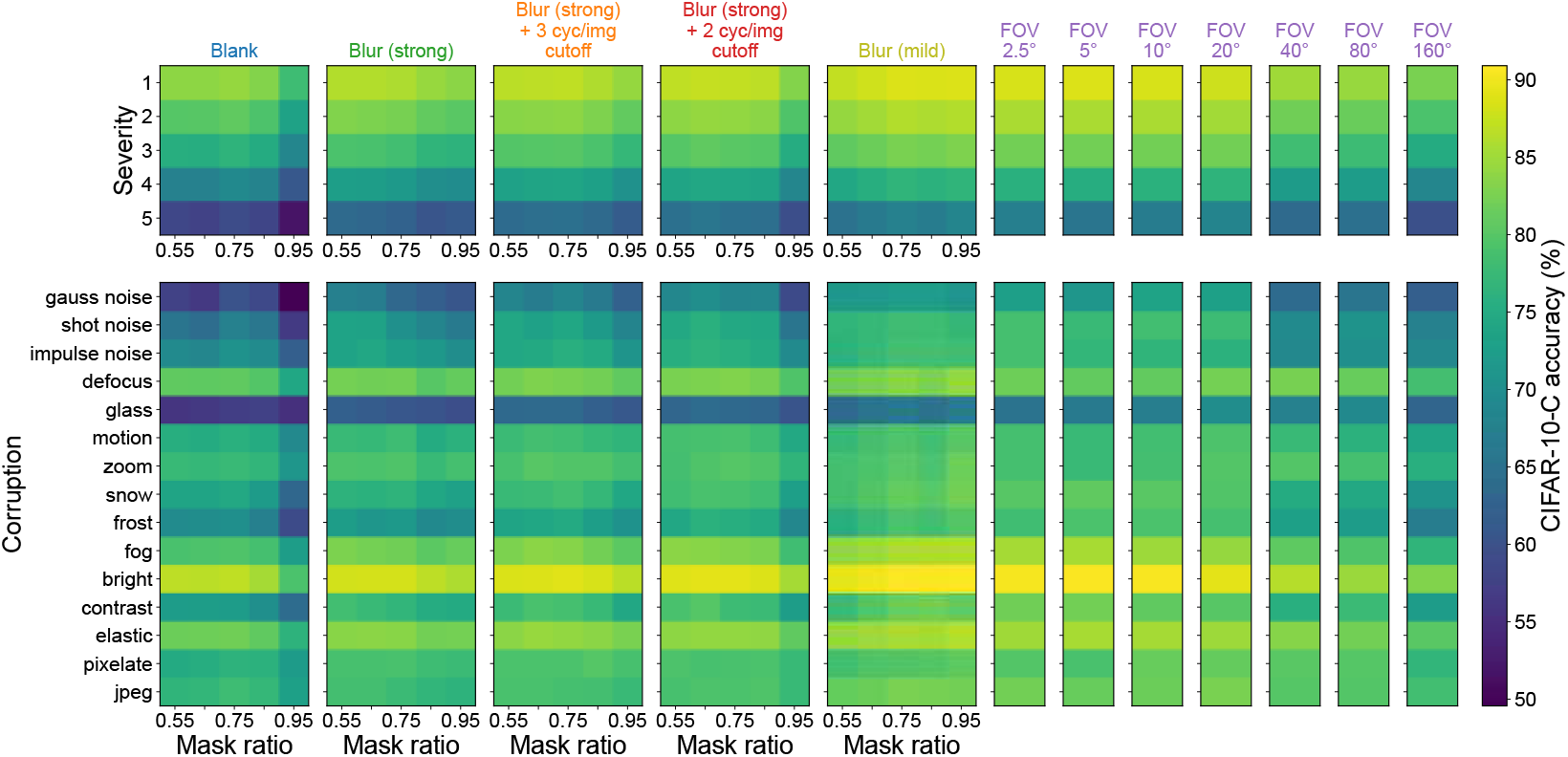
As Fig. S13 for the 50-epoch pre-training budget.

**Figure S15:**
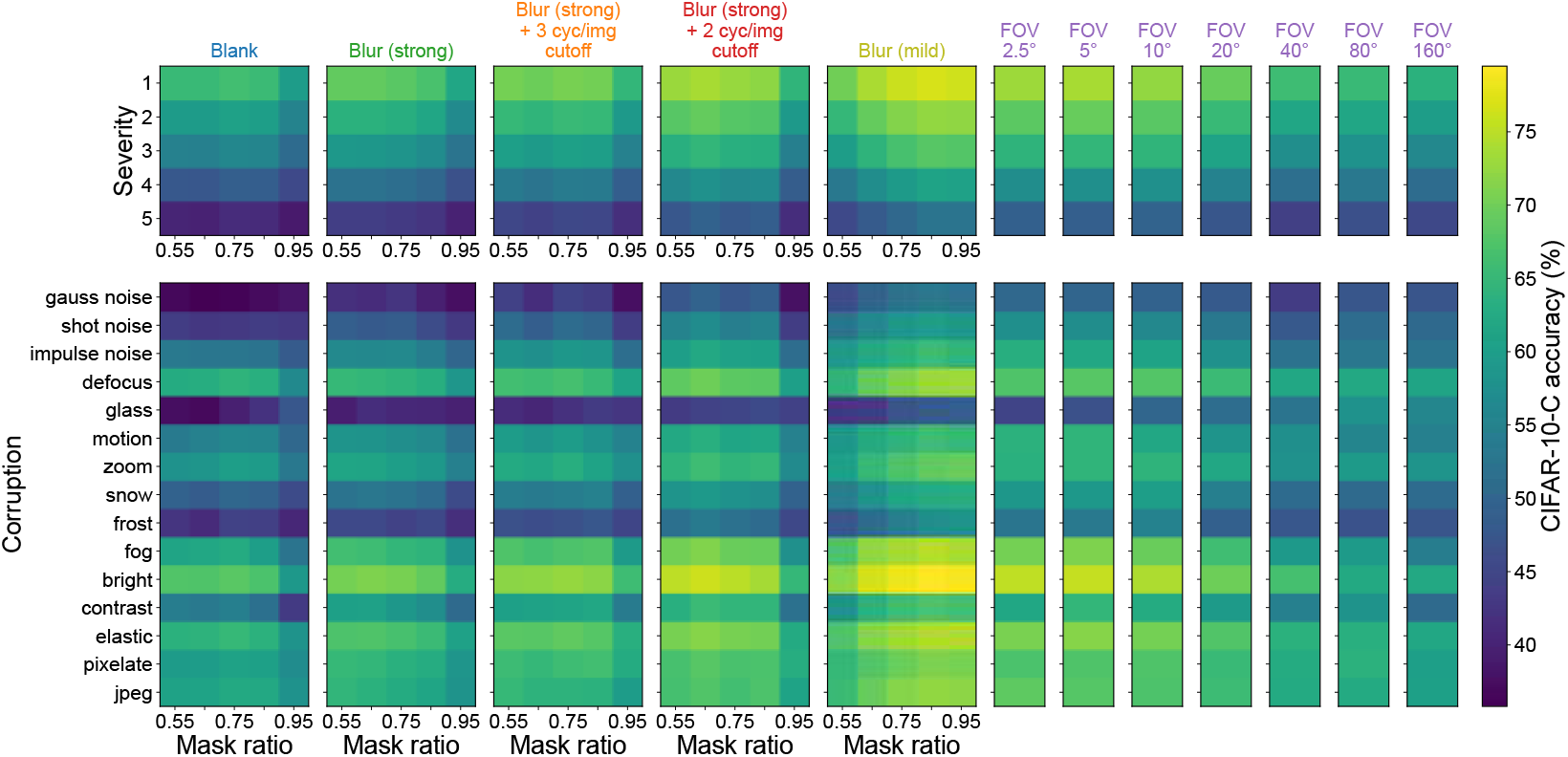
As Fig. S13 for train fraction 0.2.

**Figure S16:**
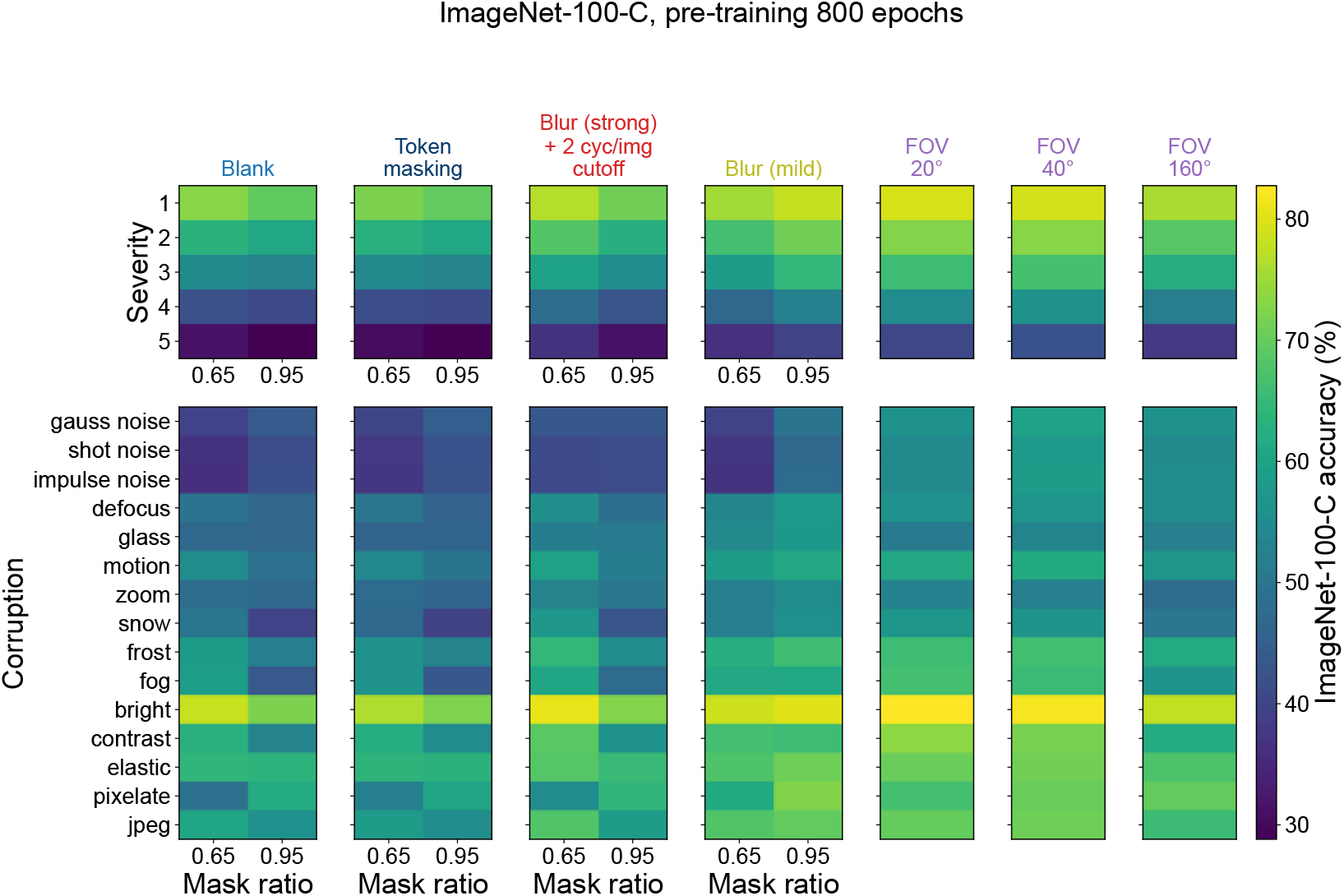
ImageNet-100-C by severity and by corruption type, full data, 800-epoch pre-training; layout as Fig. S13.

**Figure S17:**
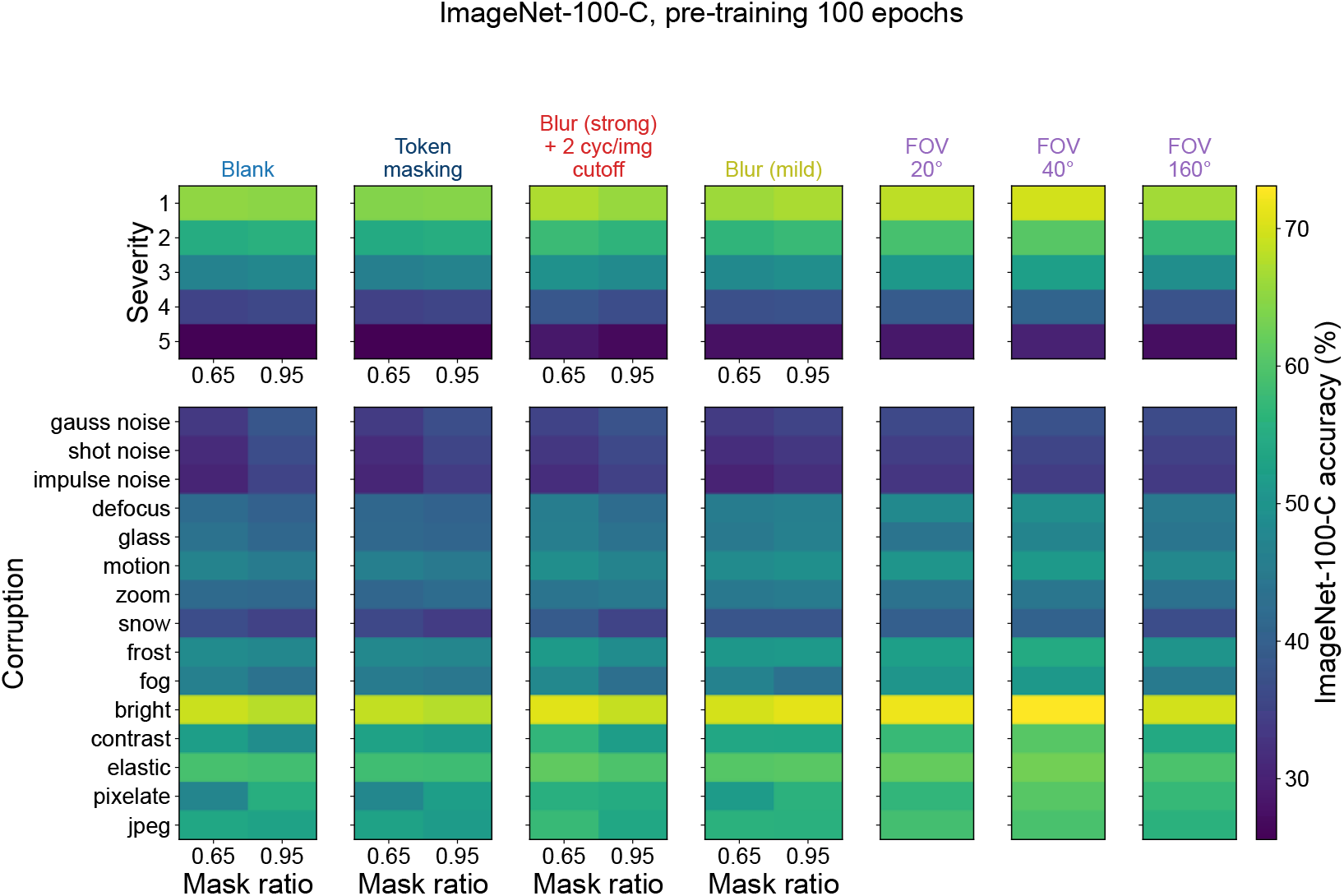
As Fig. S16 for the 100-epoch pre-training.

**Figure S18:**
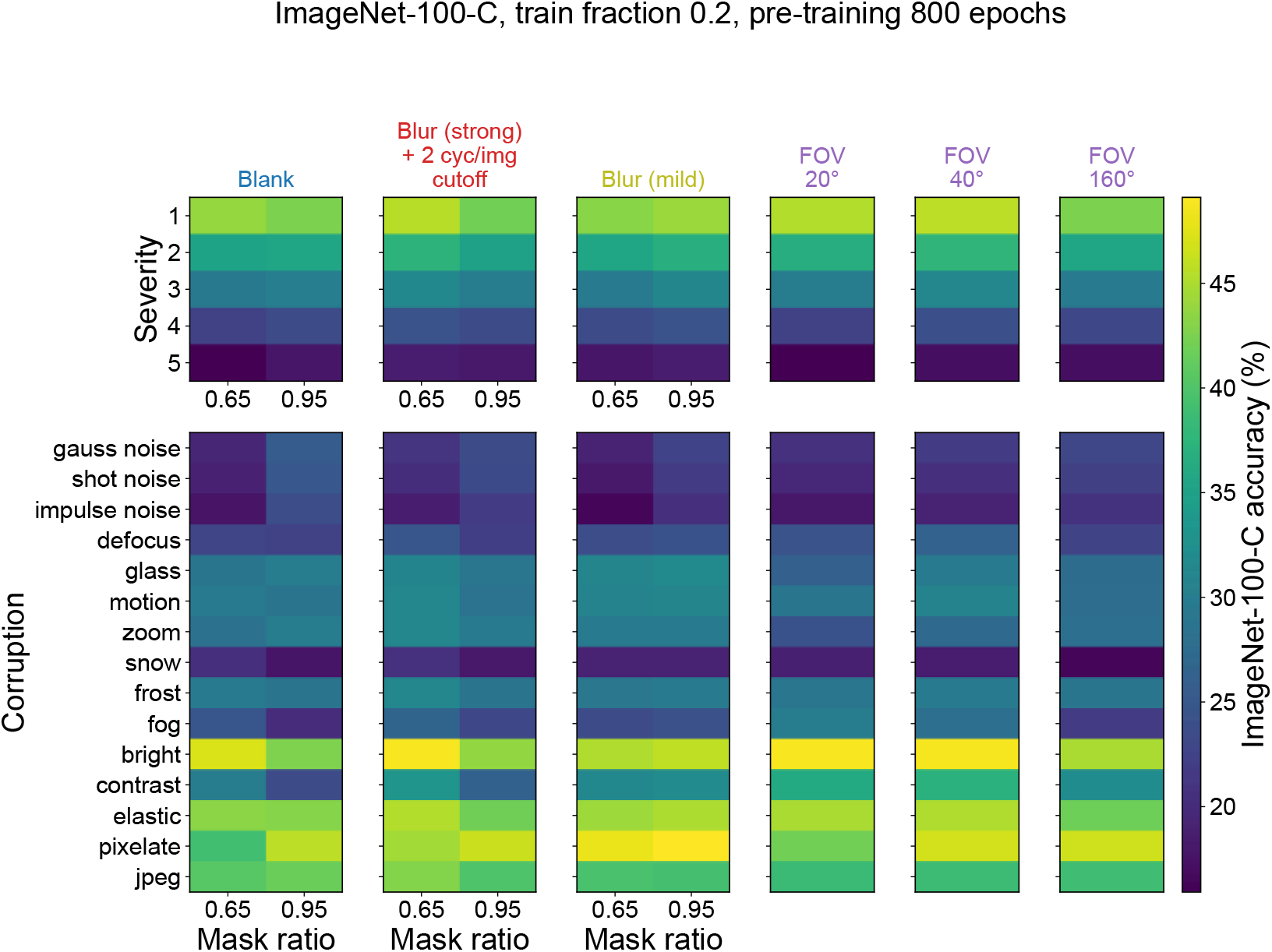
As Fig. S16 for train fraction 0.2.

**Figure S19:**
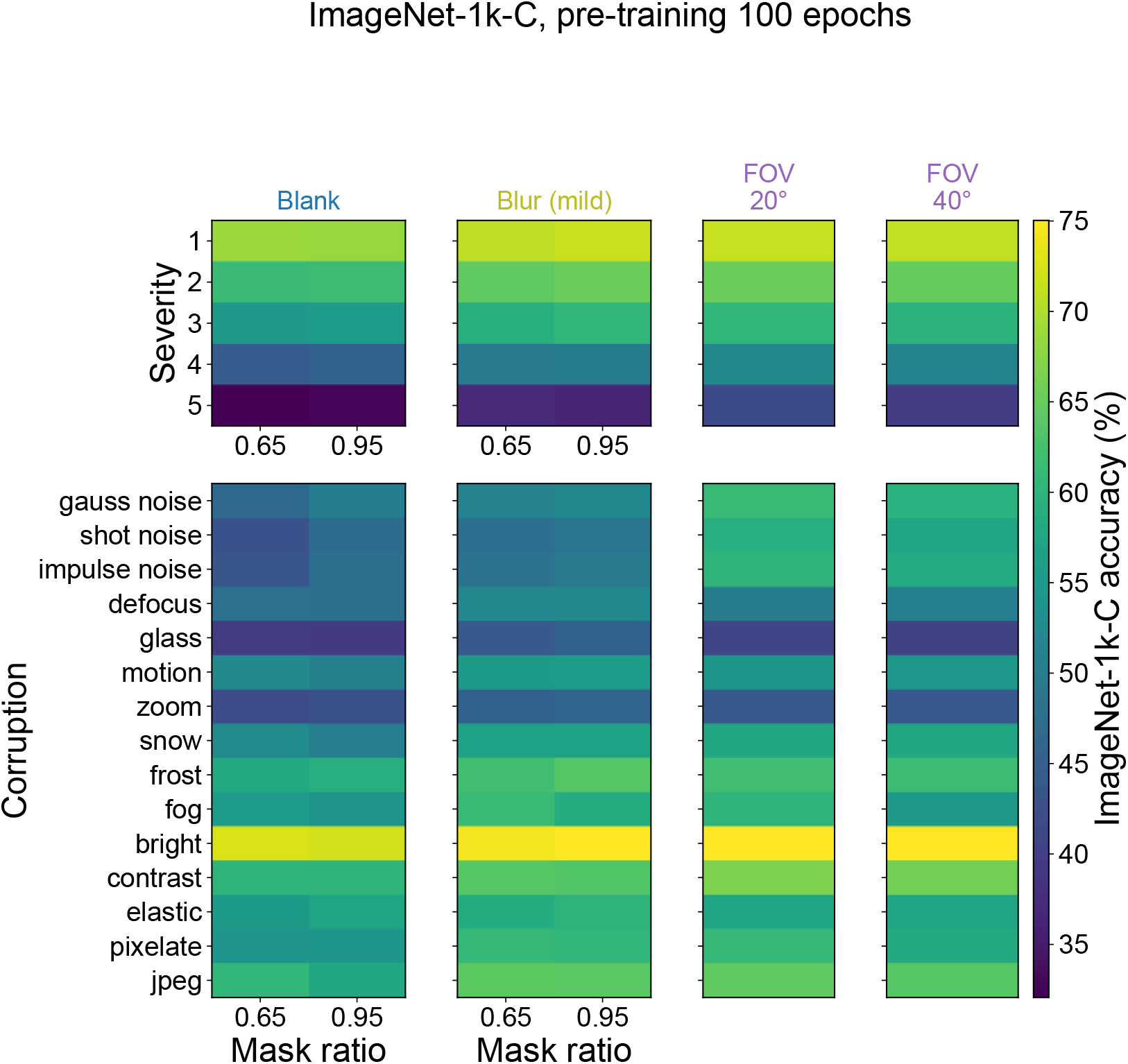
ImageNet-1k-C by severity and by corruption type; layout as Fig. S13.

**Figure S20:**
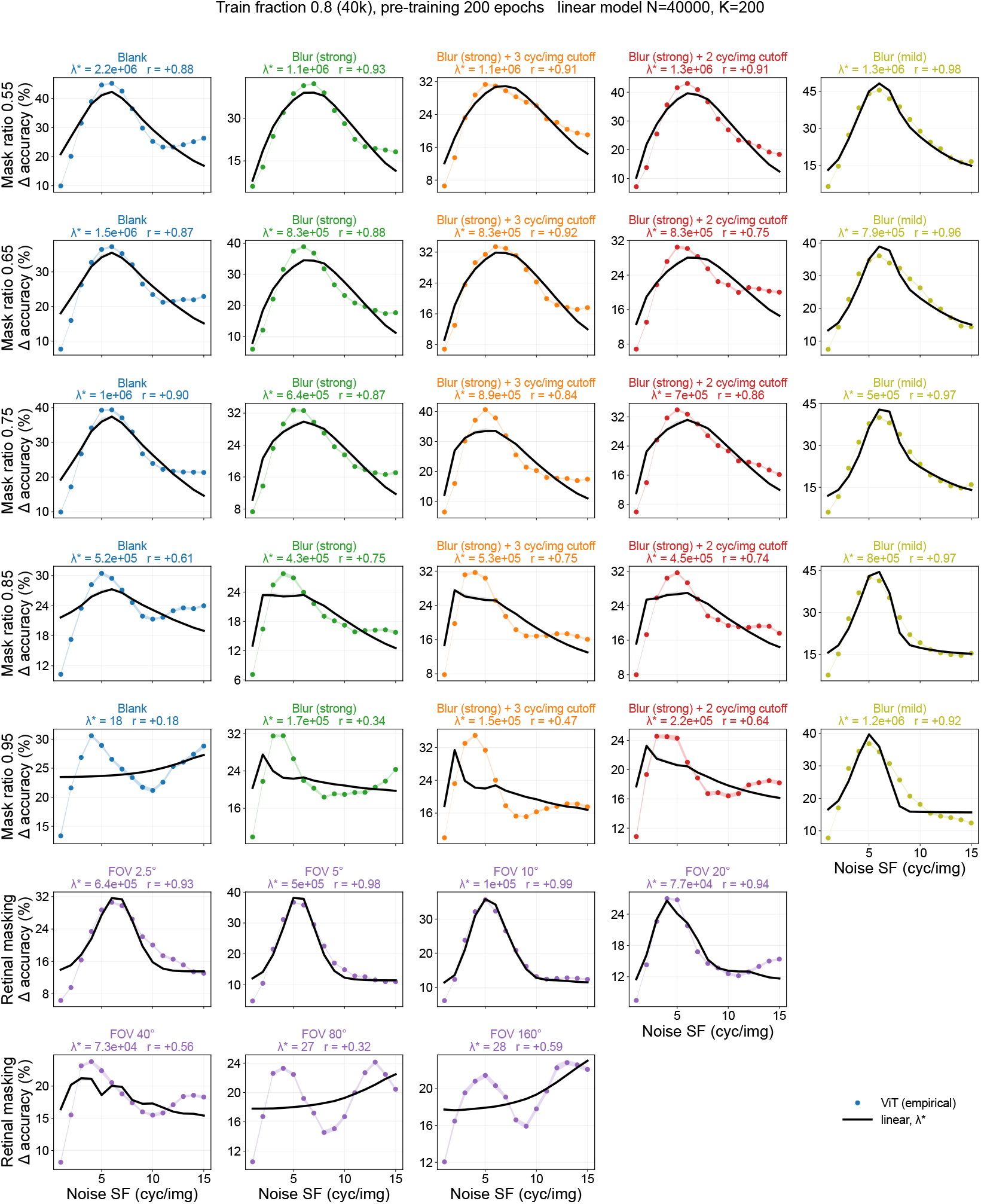
Linear-theory tuning against the ViT’s tuning, every cell of the full condition. (train fraction 0.8, 200 pre-training epochs). The ViT’s accuracy drop (points, arm colour) and the fitted linear curve (black, ± SEM over noise realisations); *λ*^∗^ and Pearson *r* in each title.

**Figure S21:**
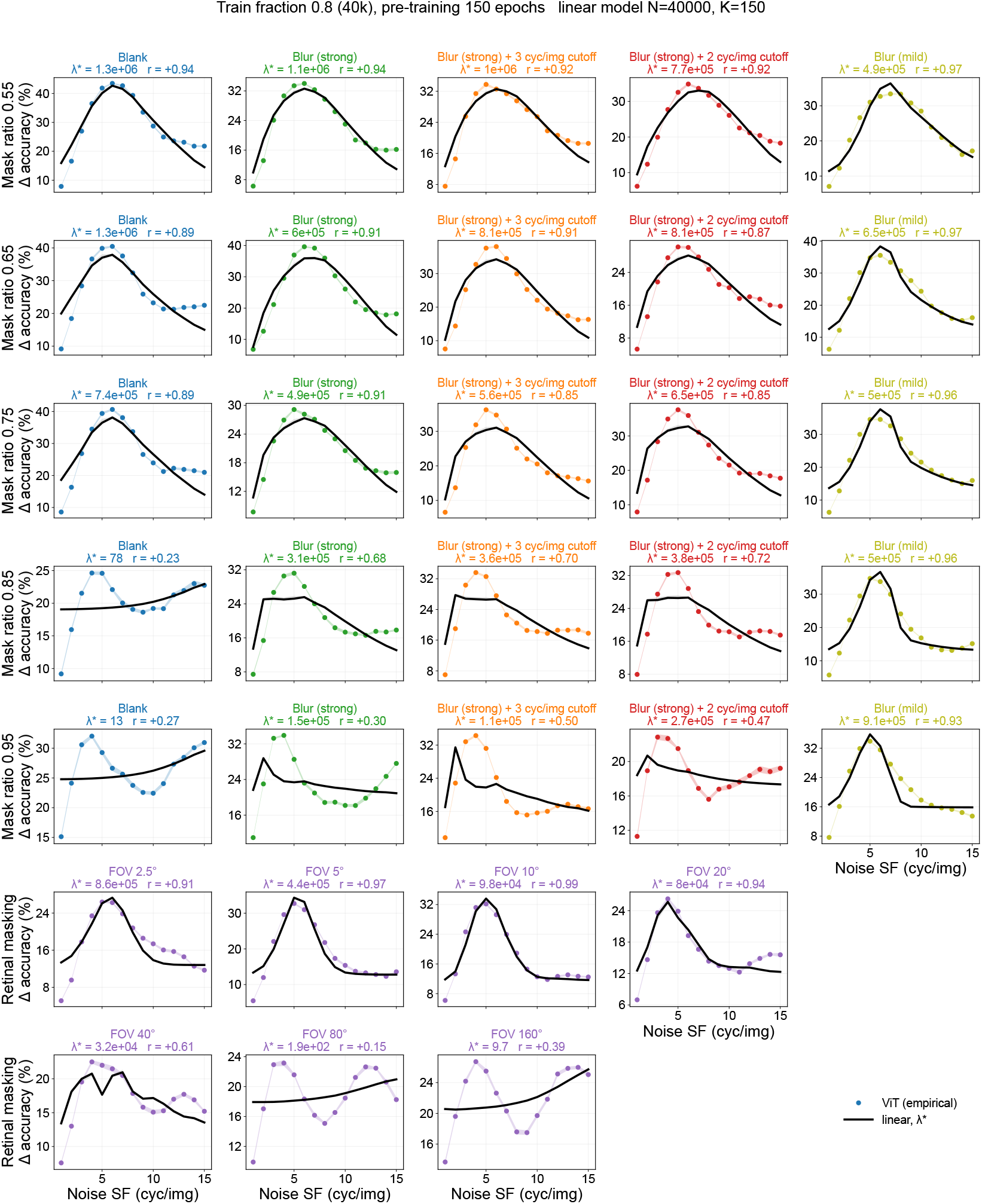
As Fig. S20 for 150 pre-training epochs.

**Figure S22:**
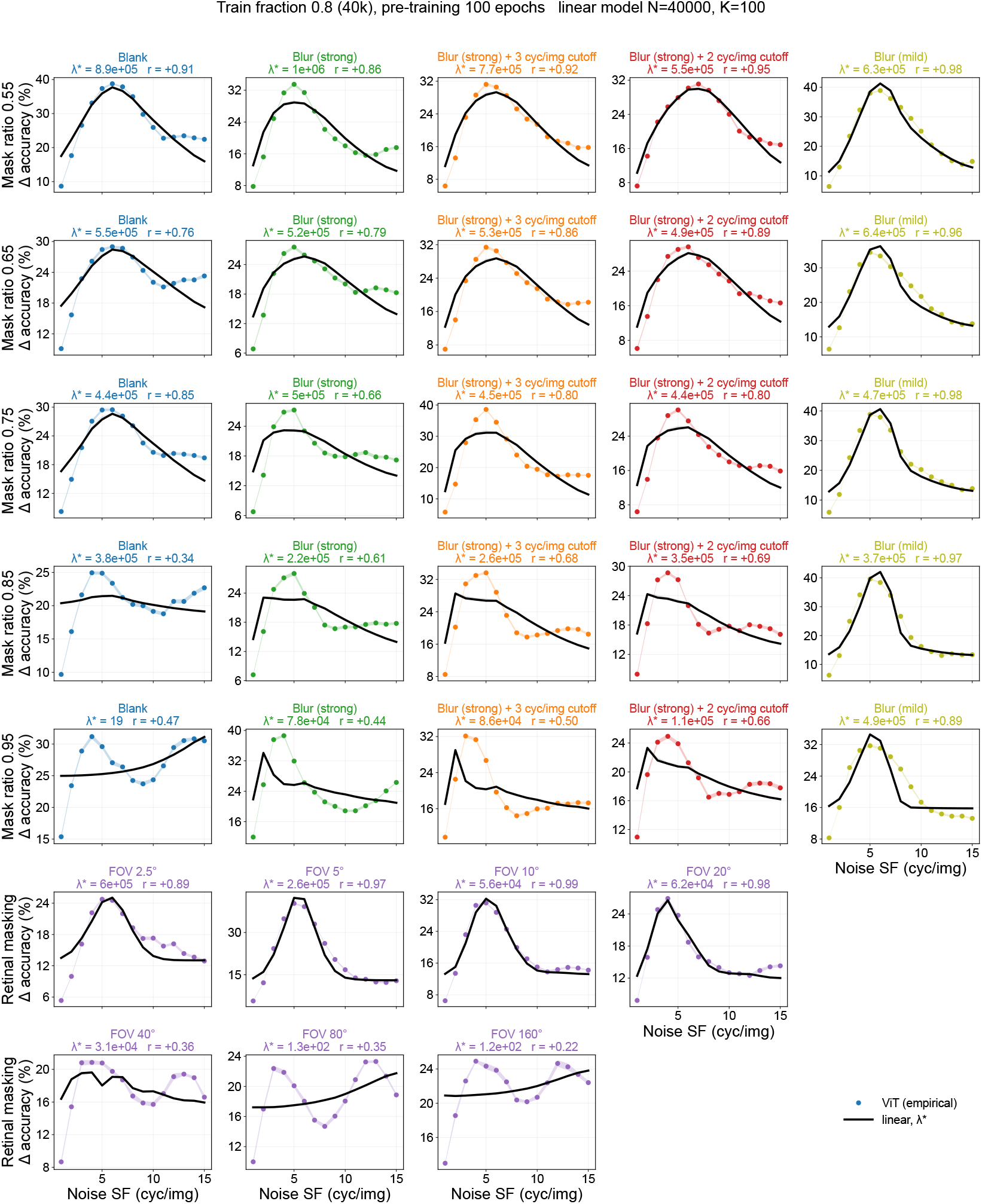
As Fig. S20 for 100 pre-training epochs.

**Figure S23:**
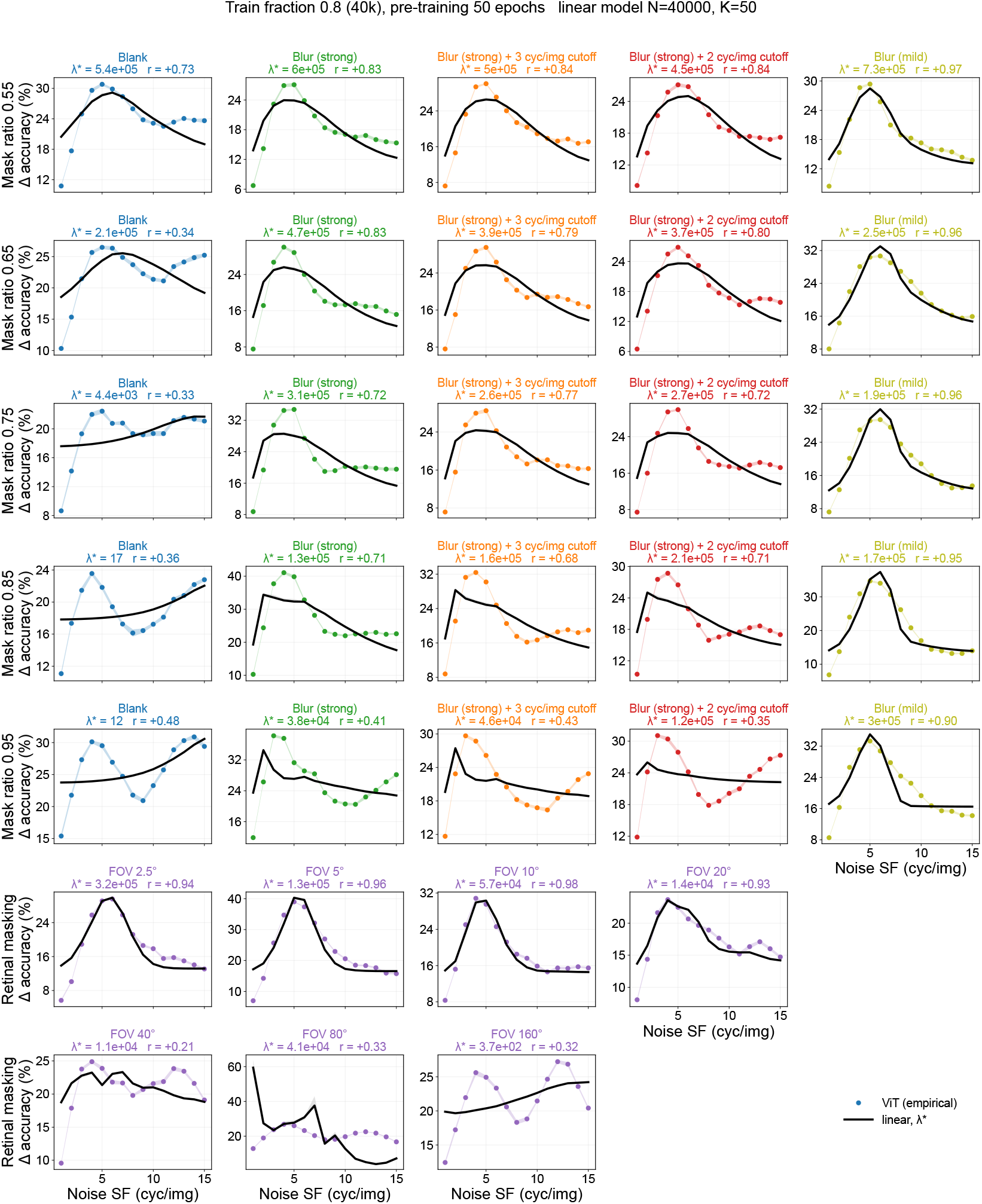
As Fig. S20 for 50 pre-training epochs.

**Figure S24:**
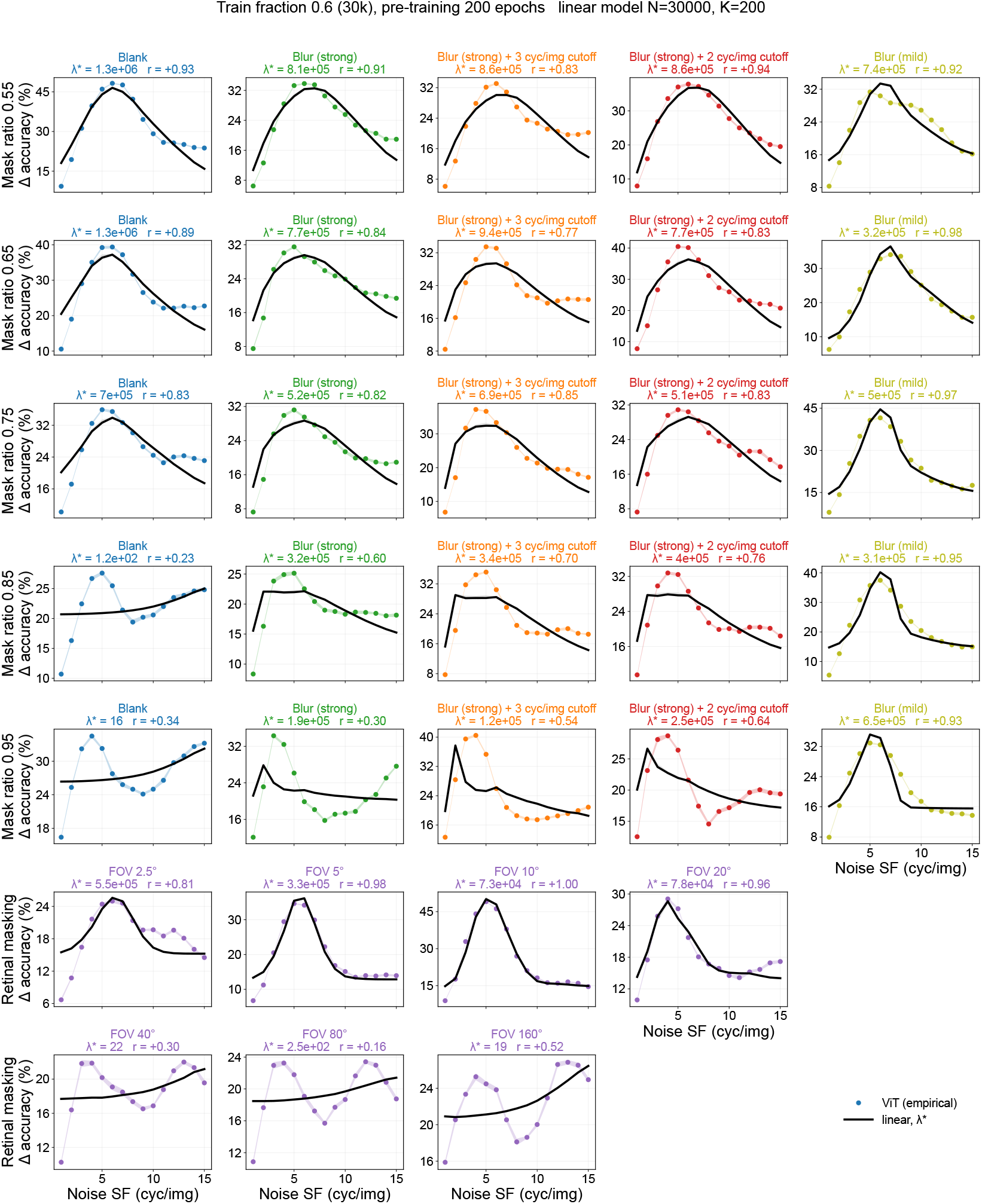
As Fig. S20 for train fraction 0.6.

**Figure S25:**
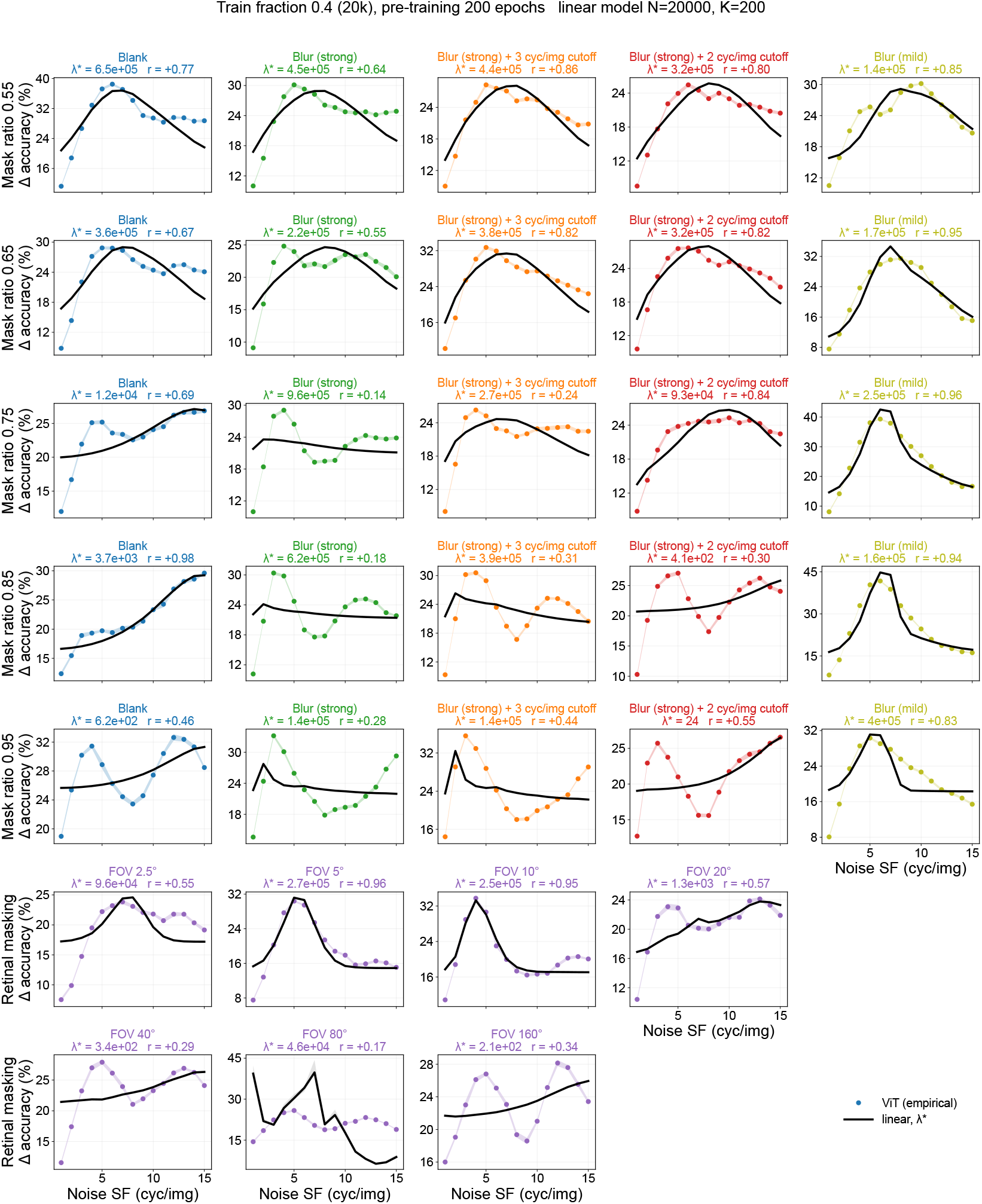
As Fig. S20 for train fraction 0.4.

**Figure S26:**
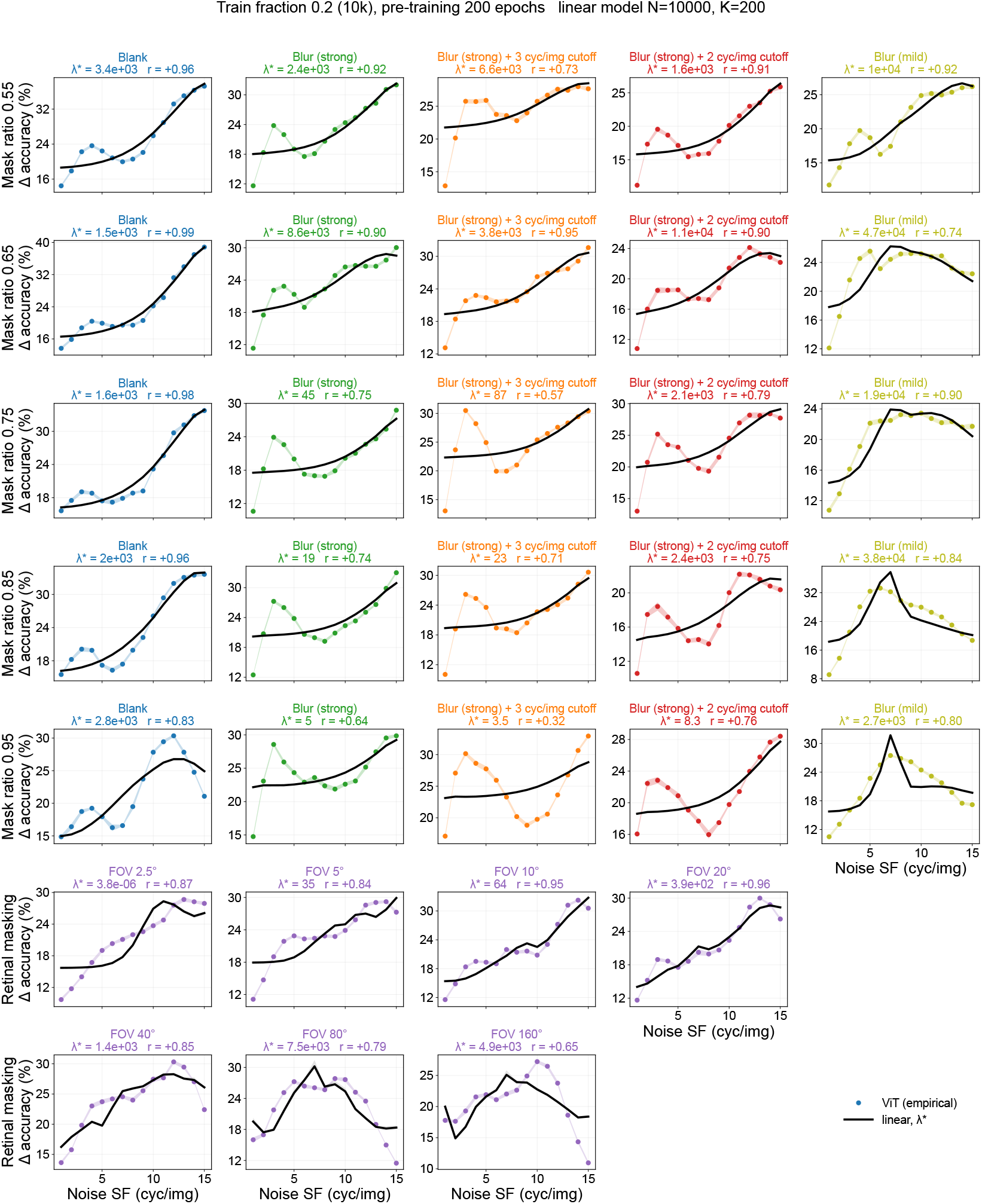
As Fig. S20 for train fraction 0.2.

**Figure S27:**
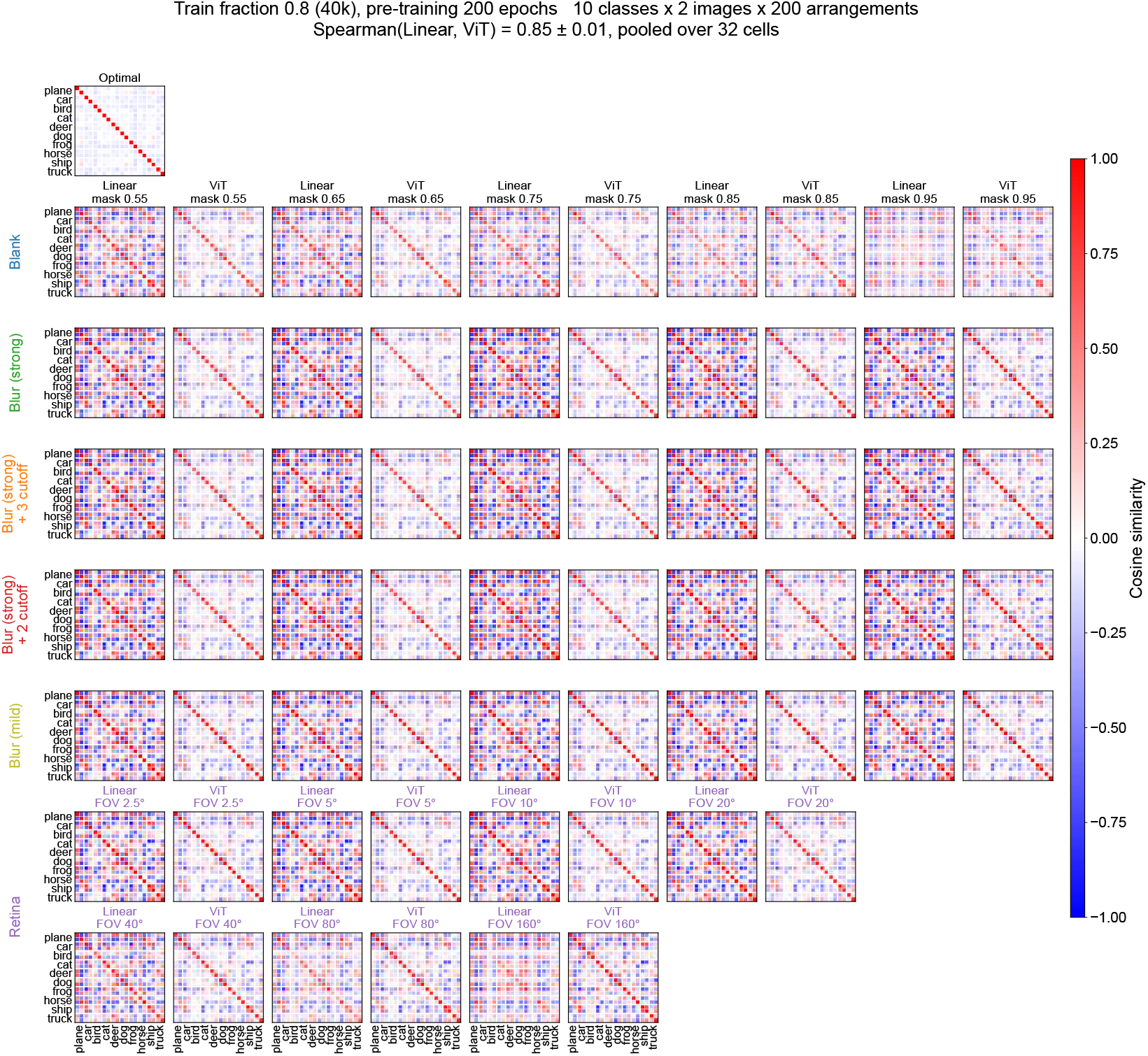
Representational similarity matrices for every corruption type in the full condition. Optimal (top), then Linear and ViT side by side for each corruption type (rows) and mask ratio (columns), with ViT-Retina (FOV 2.5–160°) in the last two rows: Each matrix is the pairwise cosine similarity between the encoder’s representations of 10 classes ×2 images ×200 corruption arrangements. The pooled Spearman correlation is given in the title.

**Figure S28:**
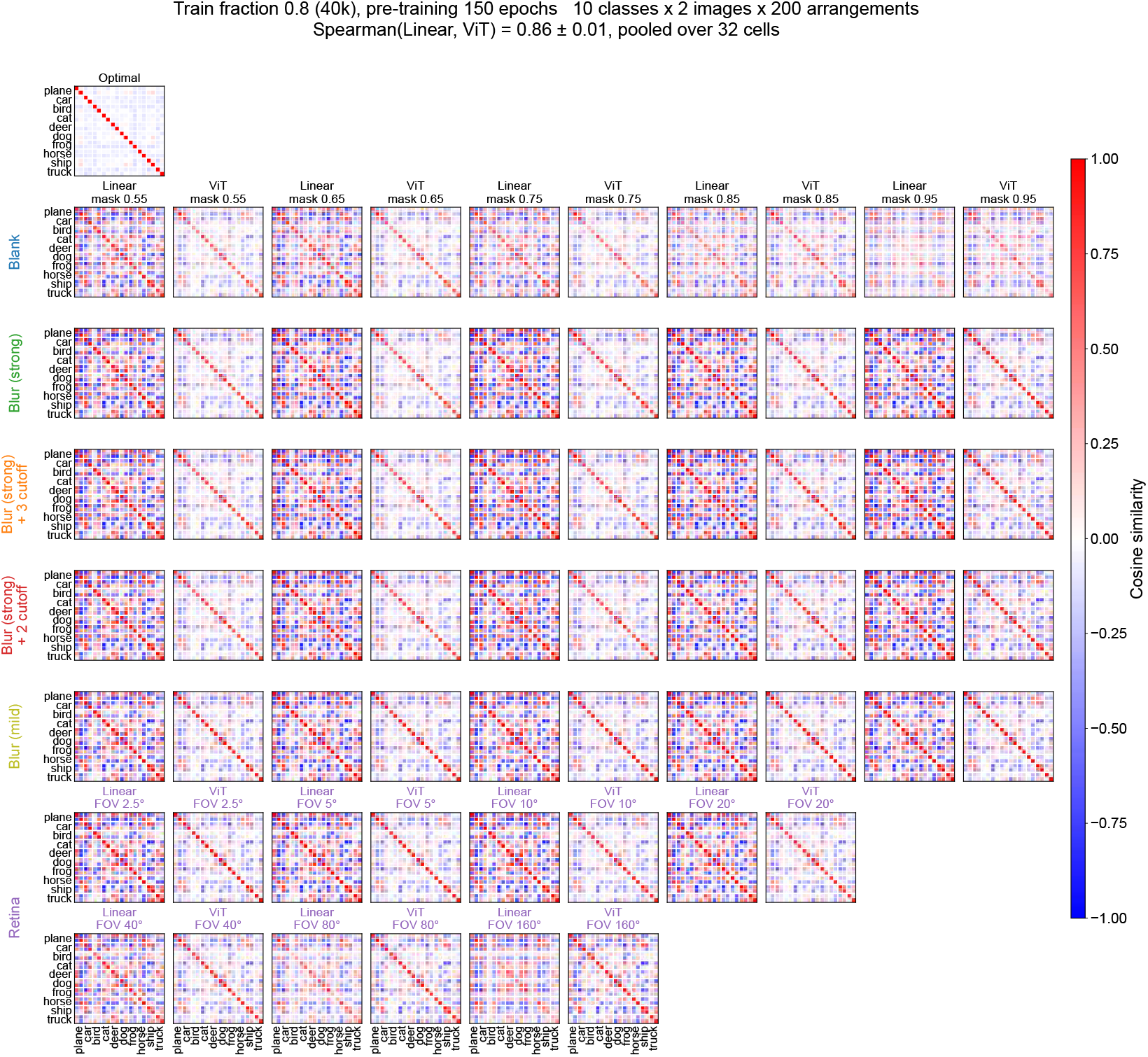
As Fig. S27 for 150 pre-training epochs.

**Figure S29:**
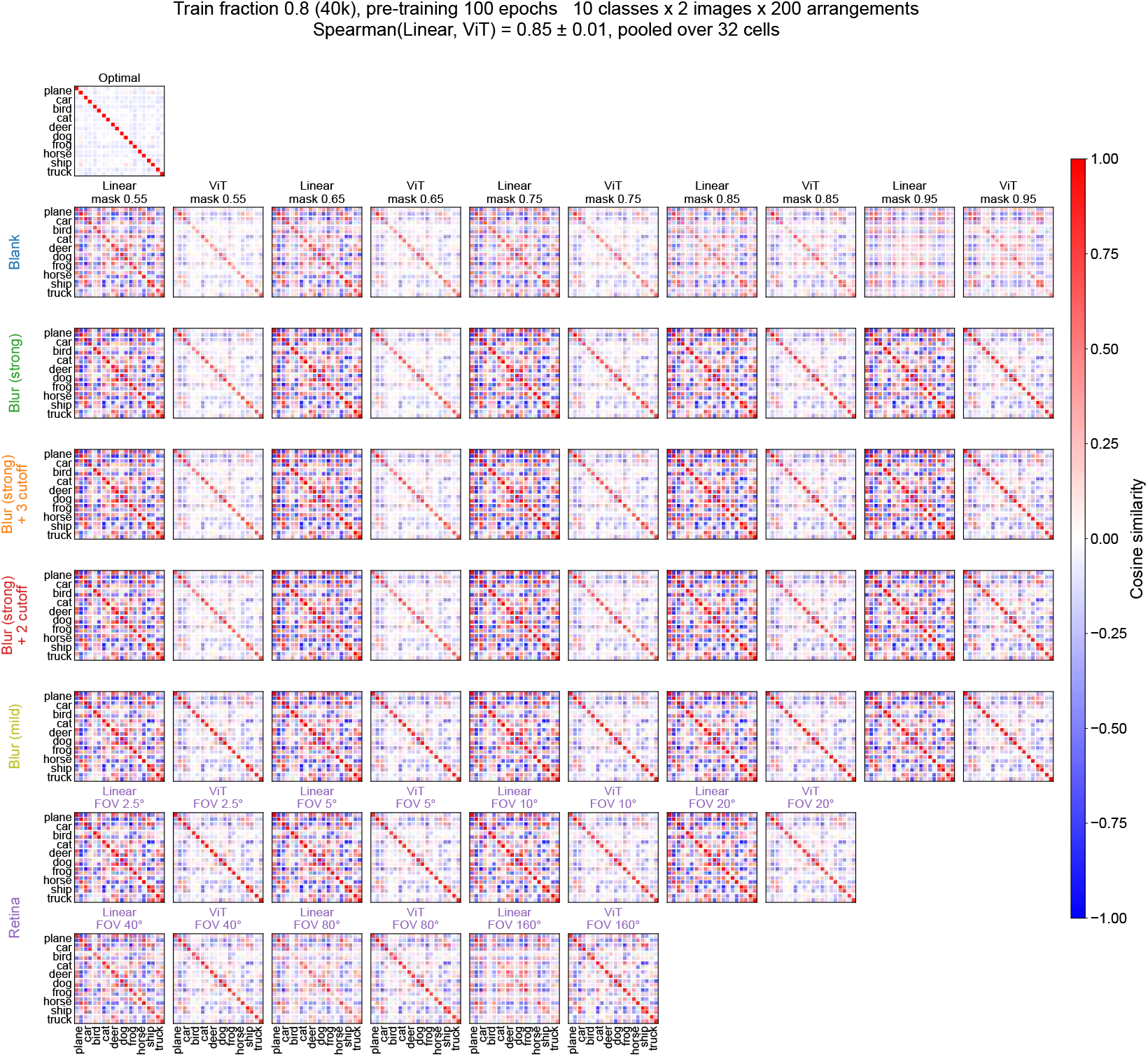
As Fig. S27 for 100 pre-training epochs.

**Figure S30:**
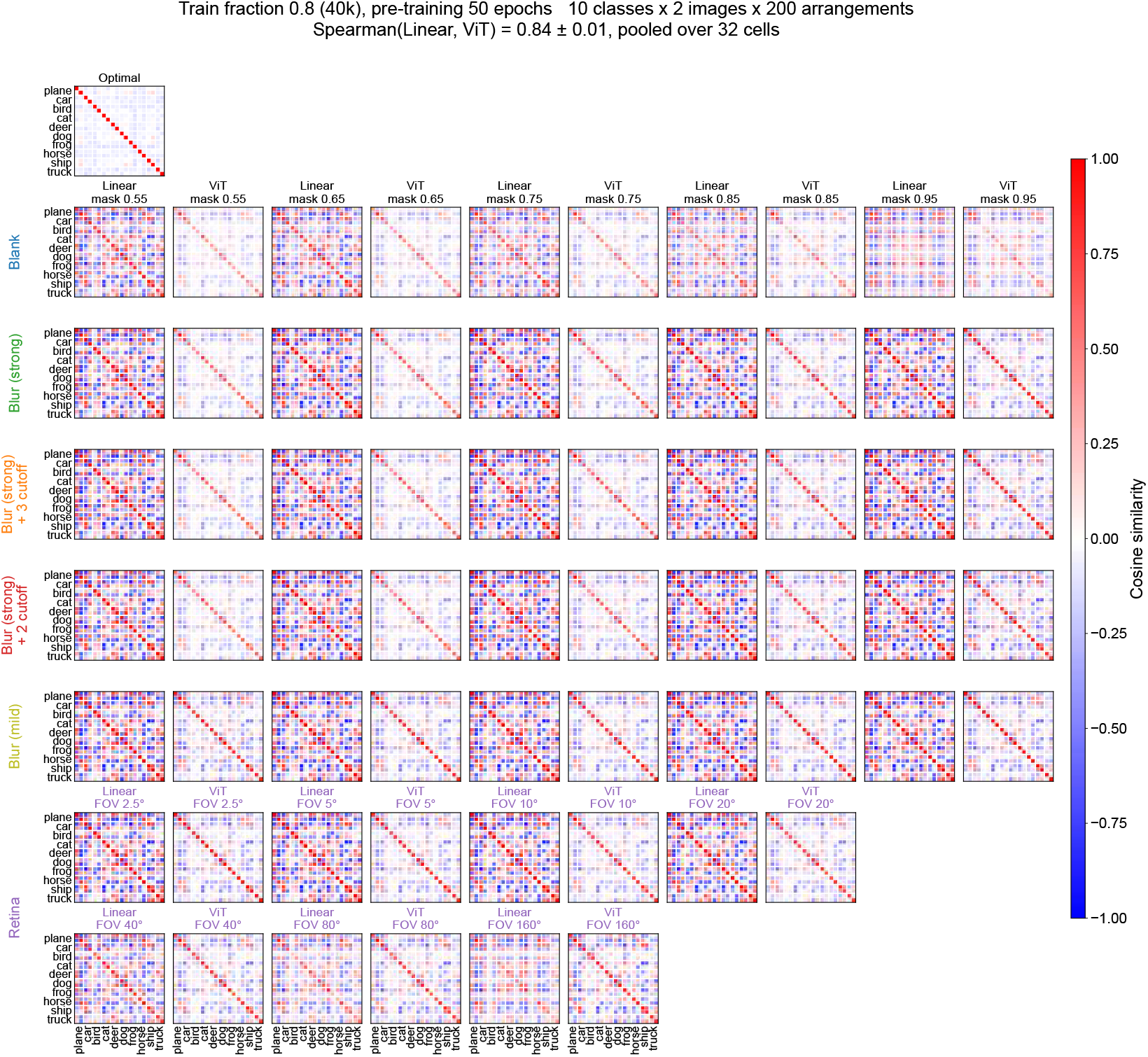
As Fig. S27 for 50 pre-training epochs.

**Figure S31:**
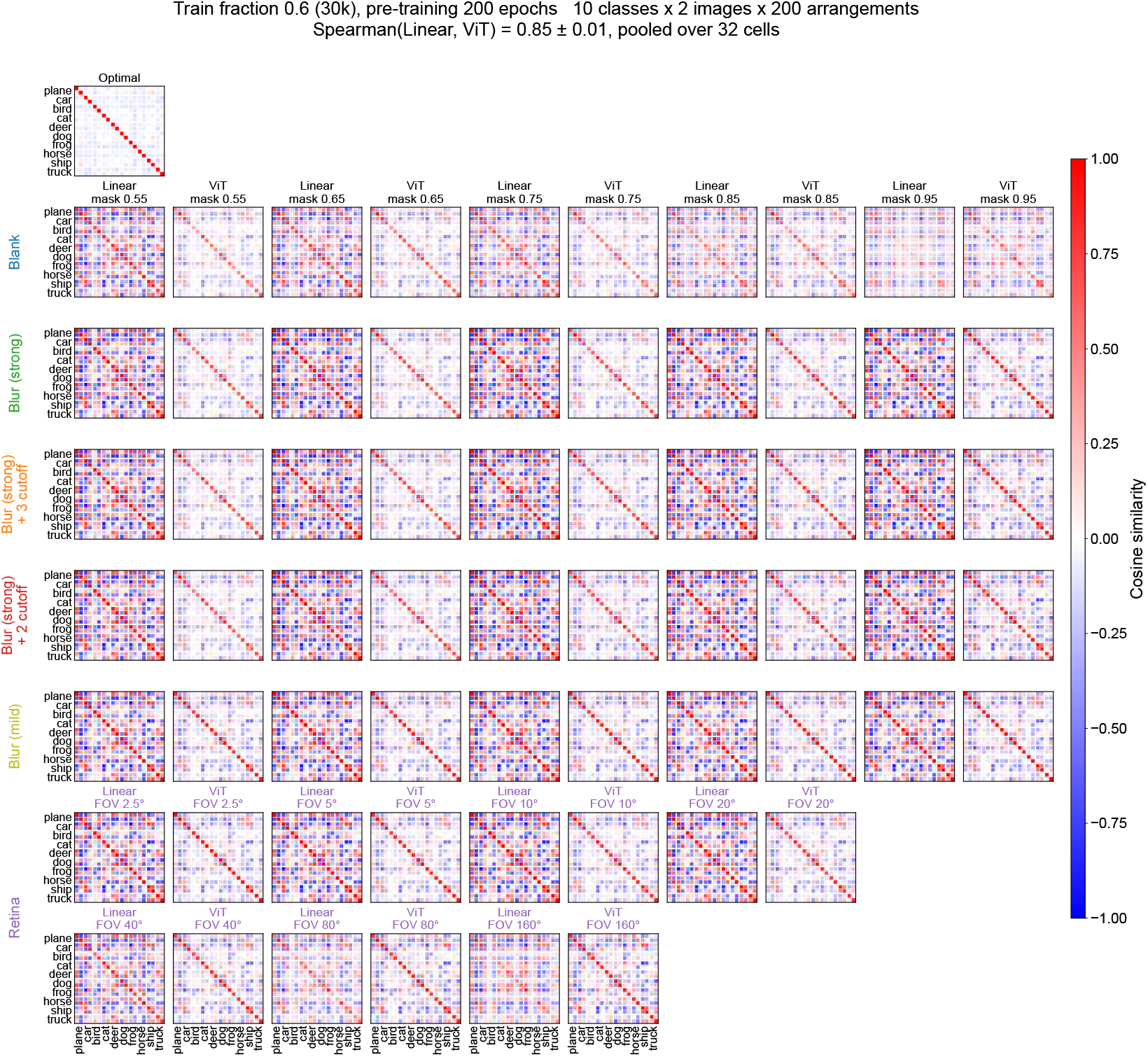
As Fig. S27 for train fraction 0.6.

**Figure S32:**
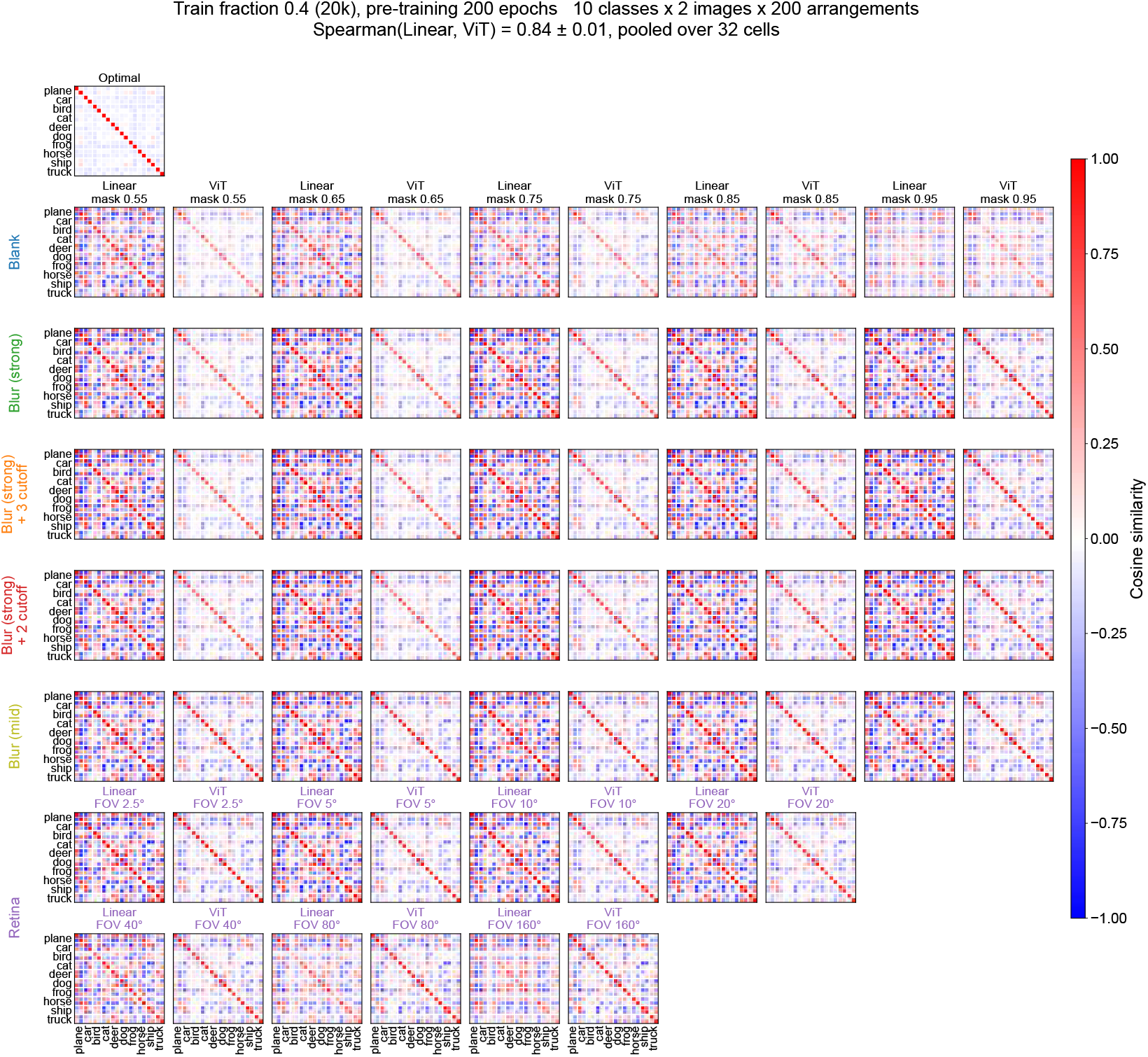
As Fig. S27 for train fraction 0.4.

**Figure S33:**
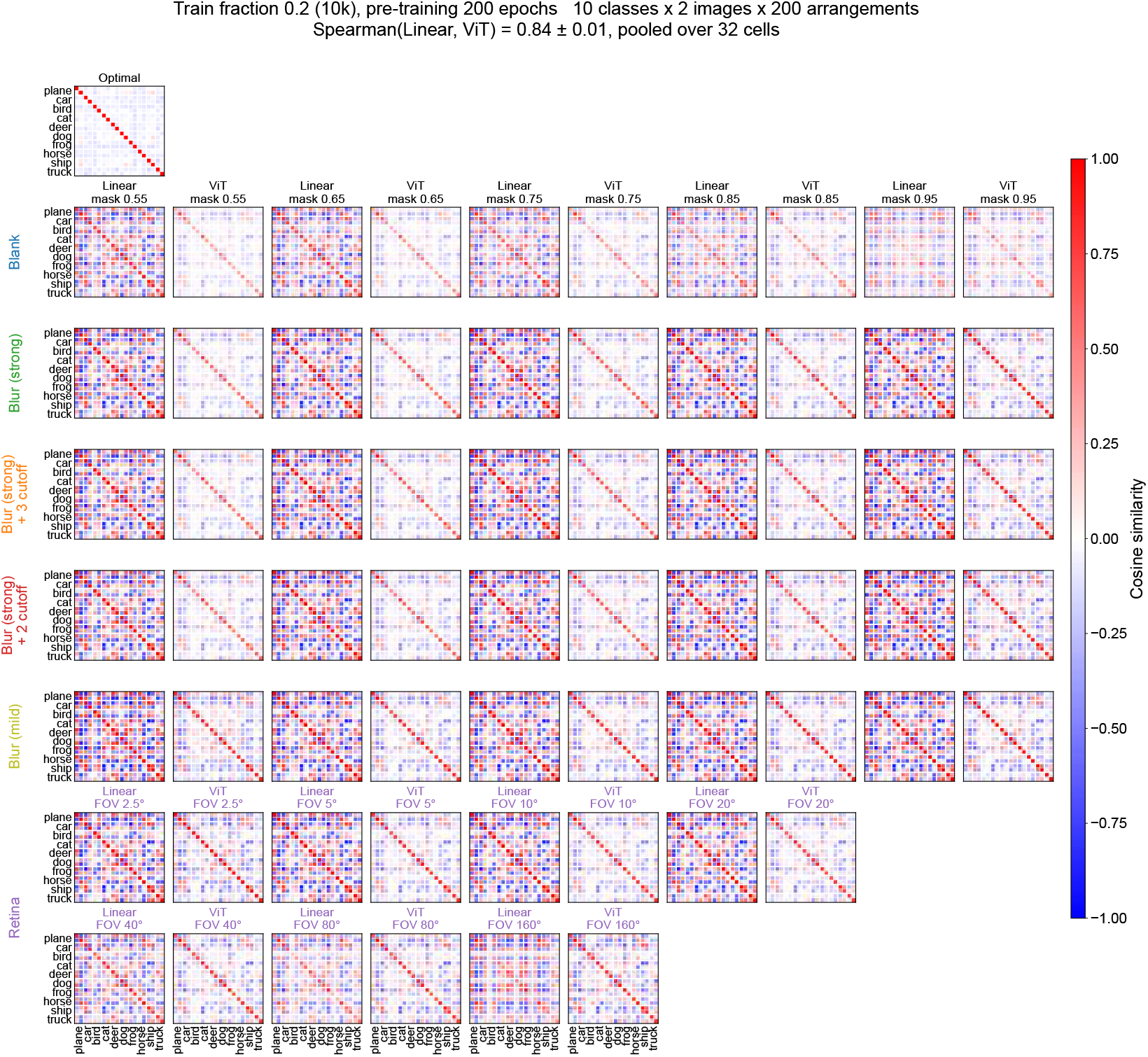
As Fig. S27 for train fraction 0.2.

**Figure S34:**
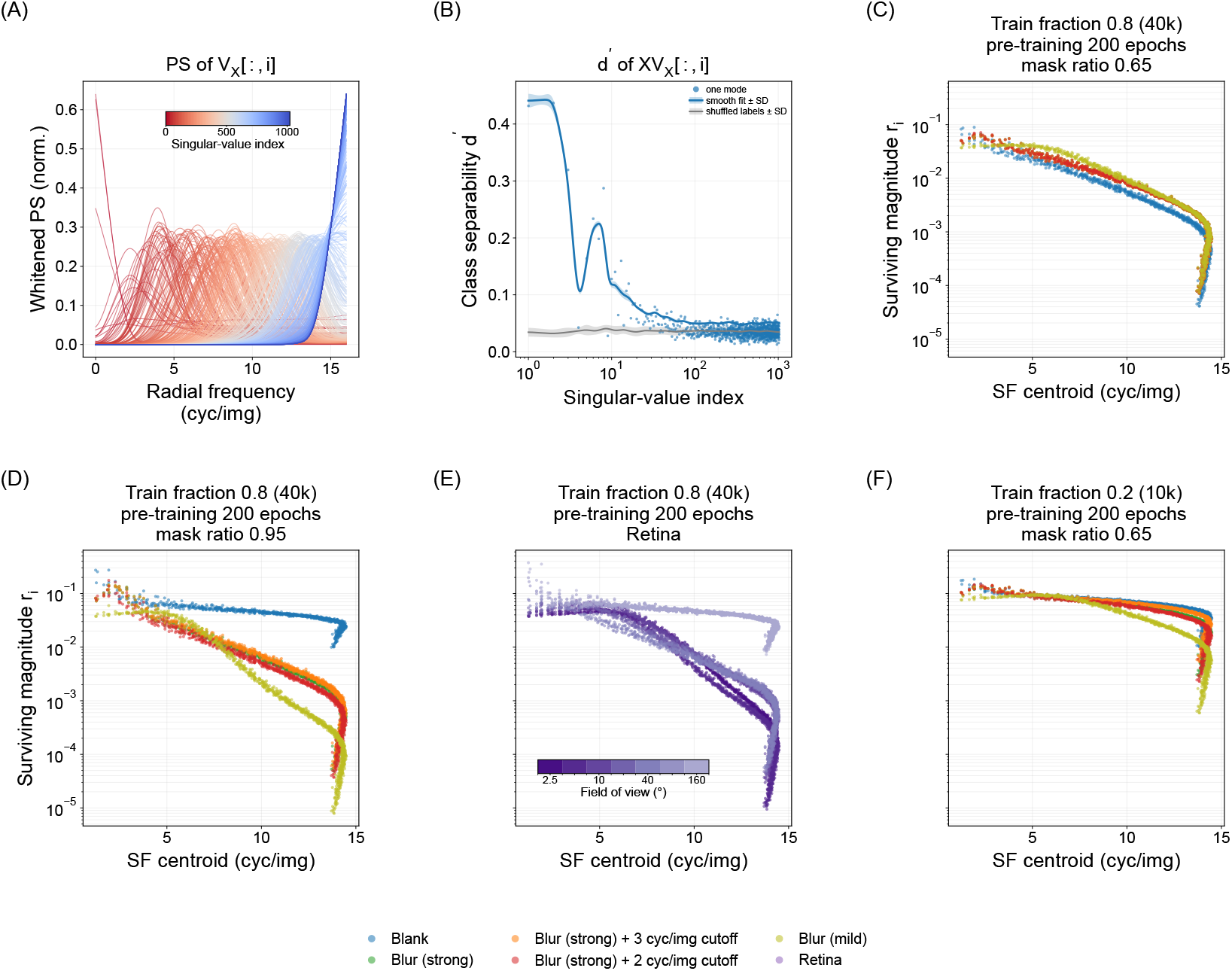
Which image modes survive the encoder. **(A)** Whitened radial power spectrum of each column of *V*_*X*_, the clean-image right singular basis, coloured by singular-value index: the higher singular-value modes are low-frequency and the lower ones high-frequency. **(B)** Class separability *d*^′^ of the projections *XV*_*X*_ [:, *i*] on held-out images (points, one per mode; smooth fit ± SD) against the shuffled-label control (grey): class information sits in the higher singular-value modes and has converged to the control by index ∼100. **(C–F)** Surviving magnitude 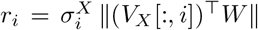 against the SF centroid of each mode. **(C)** Full condition at mask ratio 0.65: the corruption types are nearly indistinguishable. **(D)** Full condition at mask ratio 0.95: the blank masking keeps the high-SF modes while every the blur masking suppresses them by one to three orders of magnitude and keeps the class-carrying low-SF ones. **(E)** the retinal masking in the full condition, shaded by field of view: suppression is stronger at narrower FOV. **(F)** Reduced data (10K images) at mask ratio 0.65. Other mask ratios and impoverishment conditions: Figs. S35–S36.

**Figure S35:**
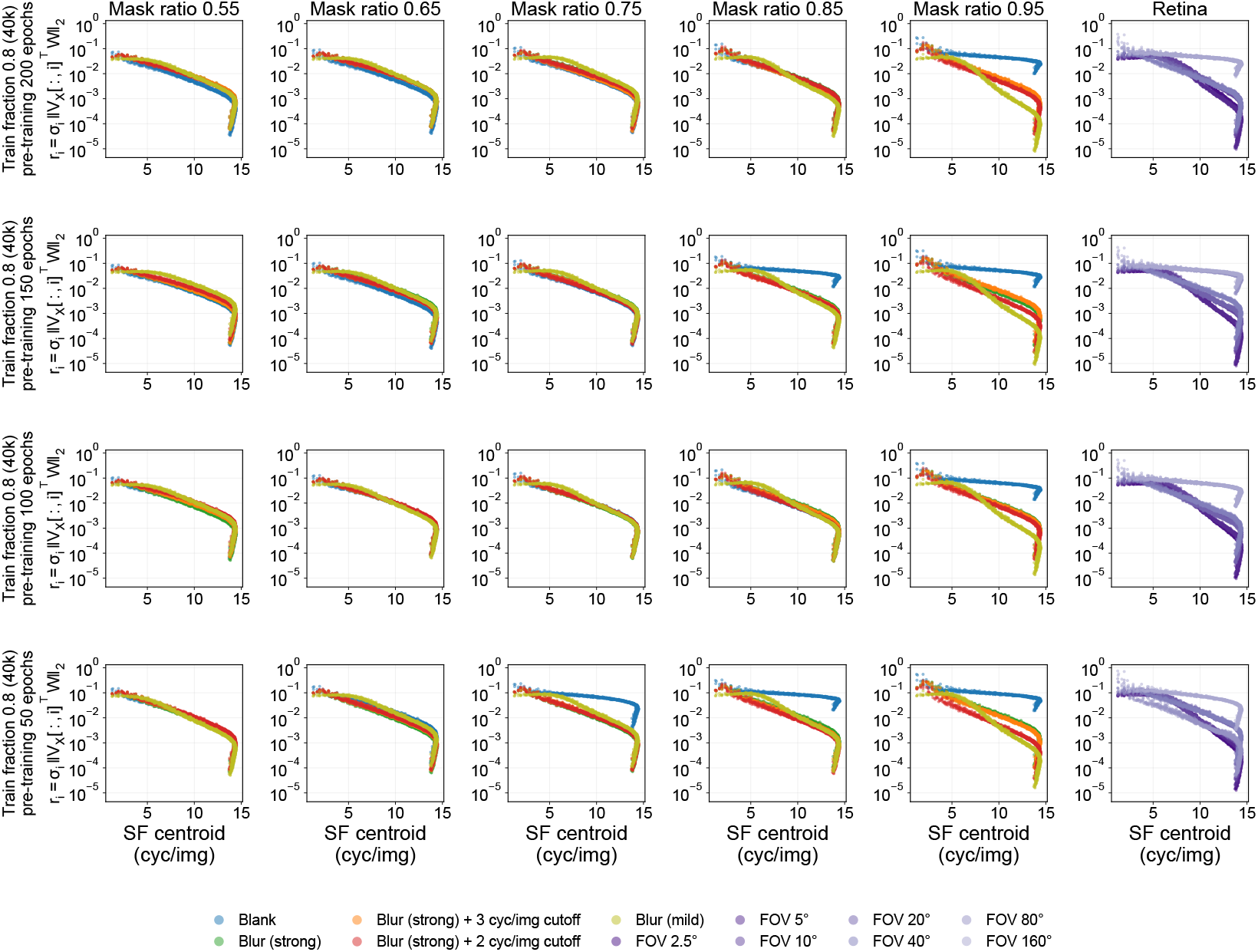
Surviving image modes along the pre-training budget axis, as Fig. S34C–F, for every mask ratio and the ViT-Retina models; one row per budget (200, 150, 100, 50 epochs).

**Figure S36:**
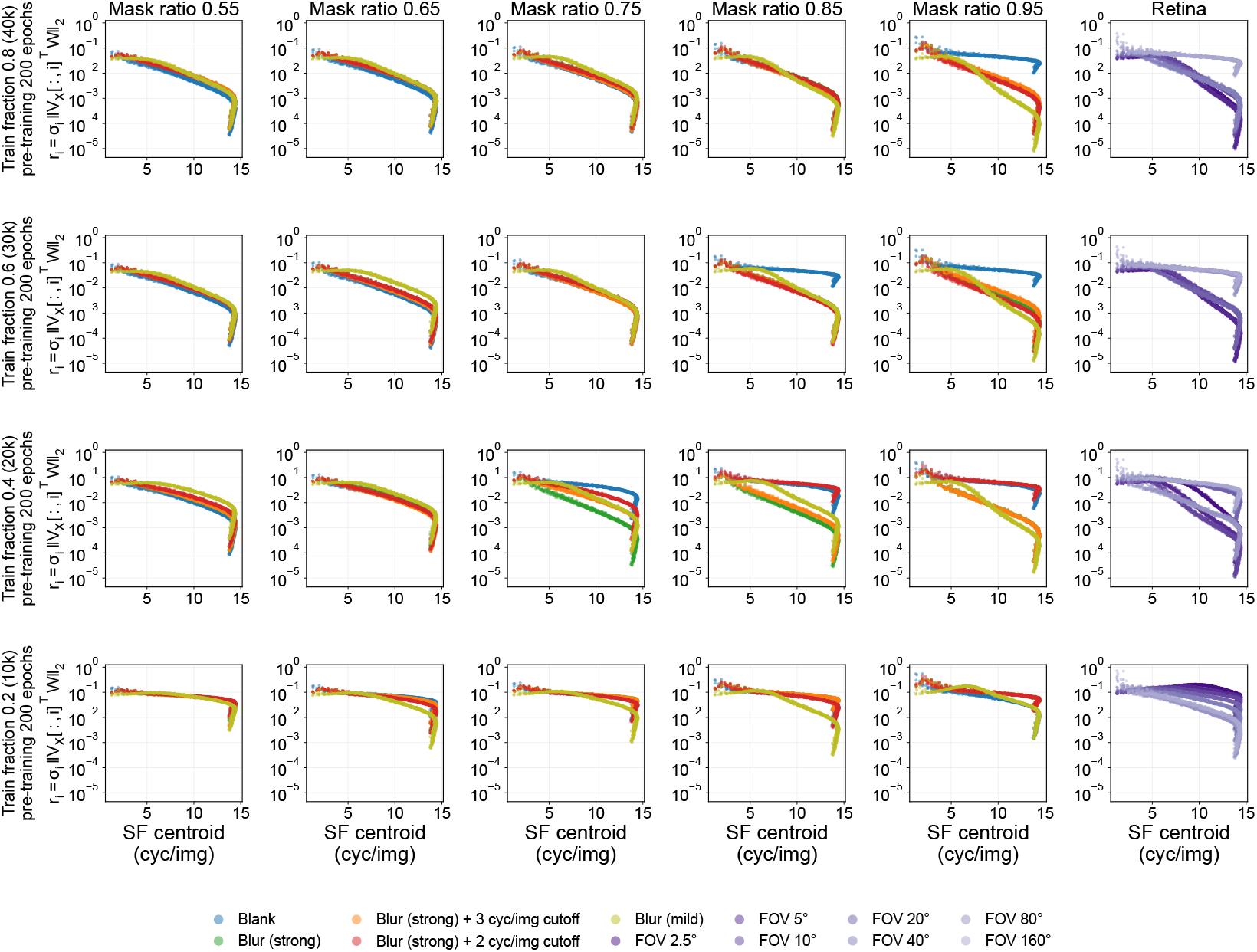
As Fig. S35 for train fractions 0.8, 0.6, 0.4 and 0.2 at 200 pre-training epochs.

**Figure S37:**
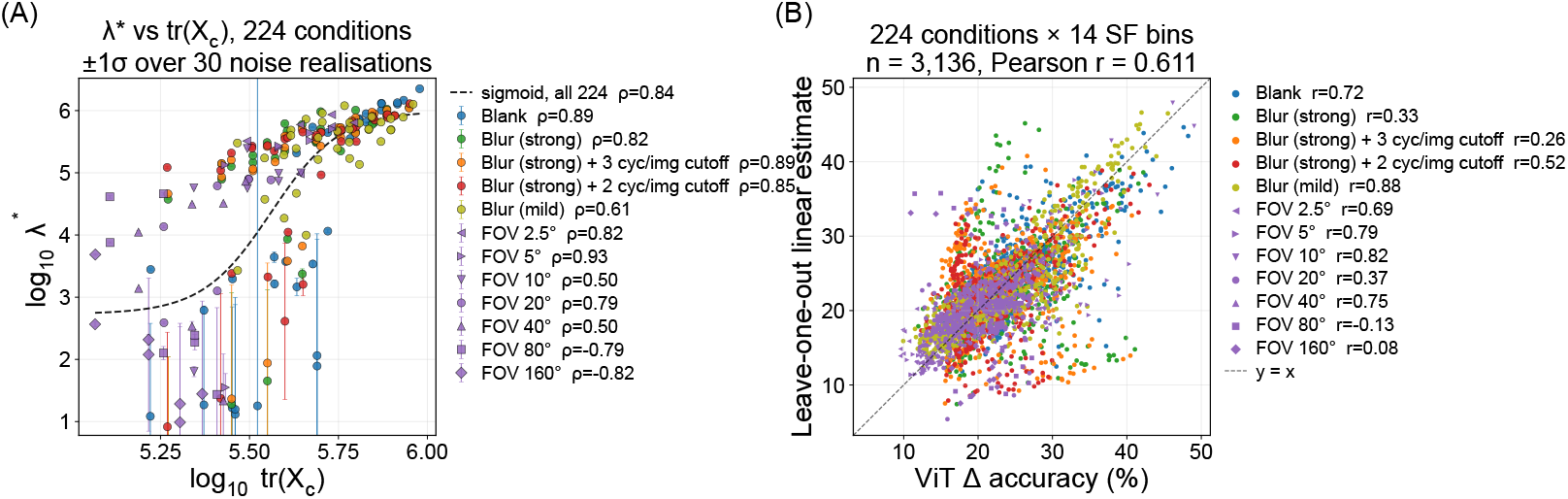
What sets. *λ*. **(A)** Fitted *λ*^∗^ against tr(*X*_*c*_) of the corrupted training set for all 224 cells (log–log; error bar 1 ± s.d. over 30 noise realisations; Spearman *ρ* = 0.84 overall, sigmoid fit dashed). **(b)** Leave-one-out: the linear tuning at the *λ* predicted from tr(*X*_*c*_) alone against the ViT’s tuning (*n* = 3,136, Pearson *r* = 0.61).

